# The evolutionarily conserved PRP4K-*CHMP4B/vps32* splicing circuit regulates autophagy

**DOI:** 10.1101/2024.05.21.595022

**Authors:** Sabateeshan Mathavarajah, Elias B. Habib, William D. Kim, Megan M. Aoki, Dale P. Corkery, Kennedy I.T. Whelan, Jordan Lukacs, Jayme Salsman, Robert J. Huber, Graham Dellaire

## Abstract

The *pre-mRNA processing factor 4 kinase (PRP4K)* is an essential gene in animal cells, making interrogation of its function challenging. Here, we report the first knockout model for PRP4K in the social amoeba *Dictyostelium discoideum*, revealing a new function in splicing events controlling autophagy. When *prp4k* knockout amoebae underwent multicellular development, we observed defects in differentiation linked to abnormal autophagy and aberrant secretion of stalk cell inducer c-di-GMP. Autophagosome-lysosome fusion was found to be impaired after PRP4K loss in both human cell lines and amoebae. Mechanistically, PRP4K loss results in mis-splicing and reduced expression of the ESCRT-III gene *CHMP4B* in human cells and its ortholog *vps32* in *Dictyostelium*, and re-expression of CHMP4B or Vps32 cDNA (respectively) restored normal autophagosome-lysosome fusion in PRP4K-deficient cells. Thus, our work reveals a novel PRP4K-*CHMP4B/vps32* splicing circuit regulating autophagy that is conserved over at least 600 million years of evolution.

## Introduction

The pre-mRNA splicing of eukaryotic genes provides functional complexity by generating multiple protein isoforms^1^. However, pathogenic transcripts that arise from mis-spliced mRNA are associated with several diseases, including transcripts of genes in the autophagy pathway^2,3^. For example, the alternative and mis-splicing of autophagy genes has been shown to impact the initiation of autophagy, the recognition of cargo, and autophagosome maturation^4–9^. Genes encoding endosomal sorting complex required for transport (ESCRT) proteins are also involved in key autophagy events such as phagophore closure and autolysosome formation^10,11^, and mis-splicing of ESCRT genes underlies defects in autophagy that contribute to the pathogenesis of cancer, eye disease and several neurological disorders^12^.

Several kinases are involved in the assembly and disassembly of spliceosome machinery, including the SRSF Protein Kinases (SRPKs), Cdc2-Like Kinases (CLKs) and pre-mRNA processing factor 4 kinase (PRP4K)^13,14^. PRP4K was the first splicing kinase identified^15,16^, is highly conserved from fission yeast to humans^14^, and regulates spliceosome assembly^17^.

Although an essential gene in most eukaryotic organisms^18–20^, partial loss of PRP4K expression is common in multiple cancers, and is associated with epithelial-to-mesenchymal transition (EMT), mitotic defects, and anoikis resistance driven by enhanced epithelial growth factor receptor (EGFR) and Yes-associated protein (YAP) signalling^21–25^. These studies point to a broad tumour suppressor role for PRP4K^20^.

RNA interference knockdown (RNAi KD) models have been the primary tool for evaluating *PRP4K* function as this gene is essential in most eukaryotic models^20^. Here, we have identified a protist that can tolerate the loss of PRP4K, *Dictyostelium discoideum*; an amoeba species that diverged from humans approximately 600 million years ago^26^. In *Dictyostelium,*

*prp4k* KO amoebae have impaired multicellular development strikingly similar to *Dictyostelium* mutants with impaired autophagy^27–29^. This led us to identify a new evolutionarily conserved role for PRP4K as a mediator of autophagosome-lysosome fusion through its kinase-dependent splicing function. Specifically, we determined that loss of *PRP4K* (or *prp4k* in amoeba) results in the mis-splicing of the ESCRT-III factor *CHMP4B* (or *vps32* in *Dictyostelium*). Thus, we have identified a novel evolutionarily conserved role for PRP4K in autophagy through regulation of *CHMP4B/vps32* pre-mRNA splicing in both human and *Dictyostelium* cells.

## Results

### Cellular localization of Dictyostelium discoideum Prp4k

The social amoeba *Dictyostelium discoideum* expresses an uncharacterized *prp4k* gene ortholog (also known as *prpf4B*), and has a unique life cycle comprised of unicellular and multicellular phases^30^. The *Dictyostelium* and human PRP4K orthologs share a highly conserved kinase domain required for splicing function (Supplementary Fig. 1). Moreover, when expressed in human cells, FLAG-tagged *Dictyostelium* Prp4k localizes to splicing speckle-like domains and co-localizes with human PRP4K^18,31^ (Fig. 1a). However, detection of endogenous *Dictyostelium* Prp4k revealed a diffusely nuclear and partly cytoplasmic localization in amoebae, which could indicate that *Dictyostelium* may lack splicing factor compartmentalization (Fig. 1b).

**Figure 1.**
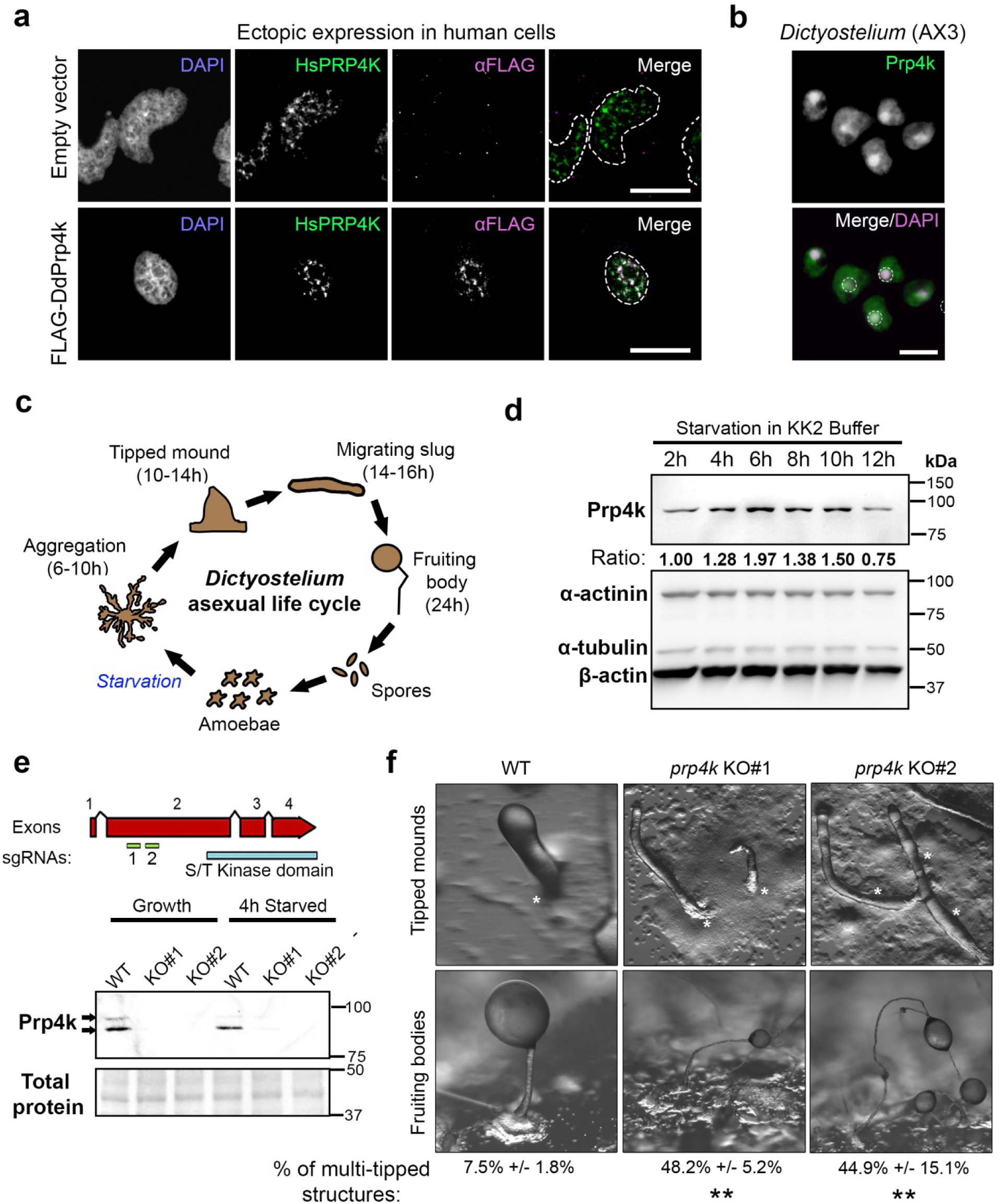
Characterization of *Dictyostelium* Prp4k localization and a multi-tip developmental phenotype in *prp4k* knockout amoebae. **(a)** Ectopic expression of FLAG-tagged *Dictyostelium* Prp4k DdPrp4k)(magenta) in HeLa cells and co-immunostaining with human PRP4K (green), which is known to localize to splicing speckles. Nuclei were counter-stained with DAPI (grey), and their outline is indicated in the merged image by white dotted lines/circles. Scale bars = 5 μm. **(b)** Immunostaining of wildtype (WT) *Dictyostelium* cells (AX3) with the custom Prp4k antibody (green). Nuclei were counterstained with DAPI (magenta), and are indicated in the merged image by dotted white circles. Scale bars = 5 μm. In a and b, the dashed white circles indicate individual nuclei. **(c)** The asexual life cycle of *Dictyostelium discoideum* consists of both a unicellular and multicellular phase that occurs over a 24 hour period. **(d)** The expression of Prp4k during starvation-induced development. Cells were starved in KK2 buffer and harvested at the indicated timepoints to assess levels of Prp4k. **(e)** Generation of a *Dictyostelium* Prp4k KO using CRISPR/Cas9. Two KO clones (derived from AX3) were isolated using two different sgRNAs that targeted the N-terminus of PRP4K (*top panel*). Prp4k protein is not present in the two *prp4k* KO amoebae. The Prp4k KO lines were validated using a custom antibody targeting the *Dictyostelium* PRP4K kinase domain (*bottom panel*). **(f)** Multicellular development of WT and *prp4k* KO lines reveals the propensity for the KO lines to form multi-tipped structures during starvation-induced development. Images were taken of tipped mounds (12 hours of development) and fruiting bodies (24 hours of development). Scale bars = 200 µm. Quantification of the multi-tipped structures (% of total structures) formed during development are shown below the images with standard deviation (n=3). Variation between groups was assessed with a one-way ANOVA, with Tukey’s post hoc analysis for pairwise comparison between groups. *p< 0.05; **p< 0.01; ***p< 0.001; ****p< 0.0001.

### Dictyostelium prp4k knockout (KO) amoebae are viable but exhibit a multi-tip developmental phenotype

Starvation triggers a developmental program marked first by cell aggregation and the formation of a multicellular structure called a mound, which then develops into a multicellular slug (Fig. 1c)^32^. Pre-stalk and pre-spore cells within the slug then terminally differentiate to form the stalk and sorus (mass of spores) of the fruiting body, which completes the life cycle^32^.

*Dictyostelium discoideum* expression data indicates that the *prp4k* gene is primarily expressed during multicellular development^33^. We examined Prp4k protein levels in *Dictyostelium* amoebae during starvation-induced aggregation. We found that Prp4k levels peak 6-10 hours post-starvation (Fig. 1d). The timing corresponds to the late aggregation and mound formation stages of the life cycle prior to when cell differentiation occurs. Therefore, we reasoned that the absence of *prp4k* at the single cell amoeba stage might be tolerated.

Using CRISPR/Cas9, we targeted the *Dictyostelium prp4k* gene at exon 2 by employing two different gRNAs (Fig. 1e). Indeed, *Dictyostelium* amoebae (AX3 strain) were able to tolerate the loss of Prp4k and we successfully generated two KO strains (KO#1 and KO#2), which were validated by reverse transcriptase quantitative PCR (RT-qPCR) and Western blot (Fig. 1e and Supplementary Fig. 2). Both KO cell lines were used in subsequent experiments. The *prp4k* KO amoebae exhibited mitotic defects that appeared to phenocopy those seen with KD of PRP4K in mammalian cells^23,34^, as indicated by the appearance of multi-nucleated amoebae (Supplementary Fig. 3a). However, this mitotic defect was most extreme in suspension-grown amoebae, and thus cells for all downstream experiments were grown strictly under adherent conditions (Supplementary Fig. 3b) ^22,34^.

When *prp4k* KO amoebae were starved, we observed abnormal tipped mounds, where many of the structures had multiple tips rather than a single tip for differentiation (Fig. 1e).

When quantified, we observed a ∼6-fold increase in multi-tipped structures in both KO strains compared to WT (Fig. 1f). However, instead of halting at multi-tipped structures, the *prp4k* KO cells complete development to form fruiting bodies with abnormal stalk morphology and reduced spore viability (Fig. 1f and Supplementary Fig. 4). The gene expression of the cell fate markers *cotC, pspA*, *ecmA*, and *ecmB* were also altered during development in *prp4k* KO amoebae; being largely suppressed compared to WT cells apart from *ecmB* which exhibited increased expression (Supplementary Fig. 5). Therefore, while *prp4k* loss can be tolerated during the growth phase of the life cycle in *Dictyostelium*, *prp4k* KO leads to severe defects in starvation-induced development.

### Dictyostelium Prp4k is required for macroautophagy and cyclic di-GMP secretion

The abnormal development of *prp4k* KO amoeba mirrors the developmental phenotypes seen in *Dictyostelium* macroautophagy mutants including *atg5^-^*, *atg6a^-^*, *atg7^-^*, *atg8^-^*, and *atg9^-^* amoebae, which all share characteristic multi-tipped mounds rather than the normal single tip mounds seen in WT development^35,36^. However, a role for PRP4K or its orthologs in macroautophagy, hereafter referred to simply as autophagy, has not been described in the literature. Therefore, we next examined autophagy in *prp4K KO* amoebae during starvation. One method of examining autophagy in *Dictyostelium* is to measure the turnover of cytoplasmic proteins during starvation^37^. For example, autophagy mutants accumulate electron-dense material in the cytoplasm that can be visualized by transmission electron microscopy (TEM). When we examined WT and *prp4K KO* amoebae grown for 36 hours in media lacking amino acids (FM medium) to induce autophagy, the cytoplasm was significantly more electron dense in the *prp4k* KO cells compared to WT cells (Fig. 2b), which is consistent with impaired autophagy after Prp4k loss (Fig. 2b).

**Figure 2.**
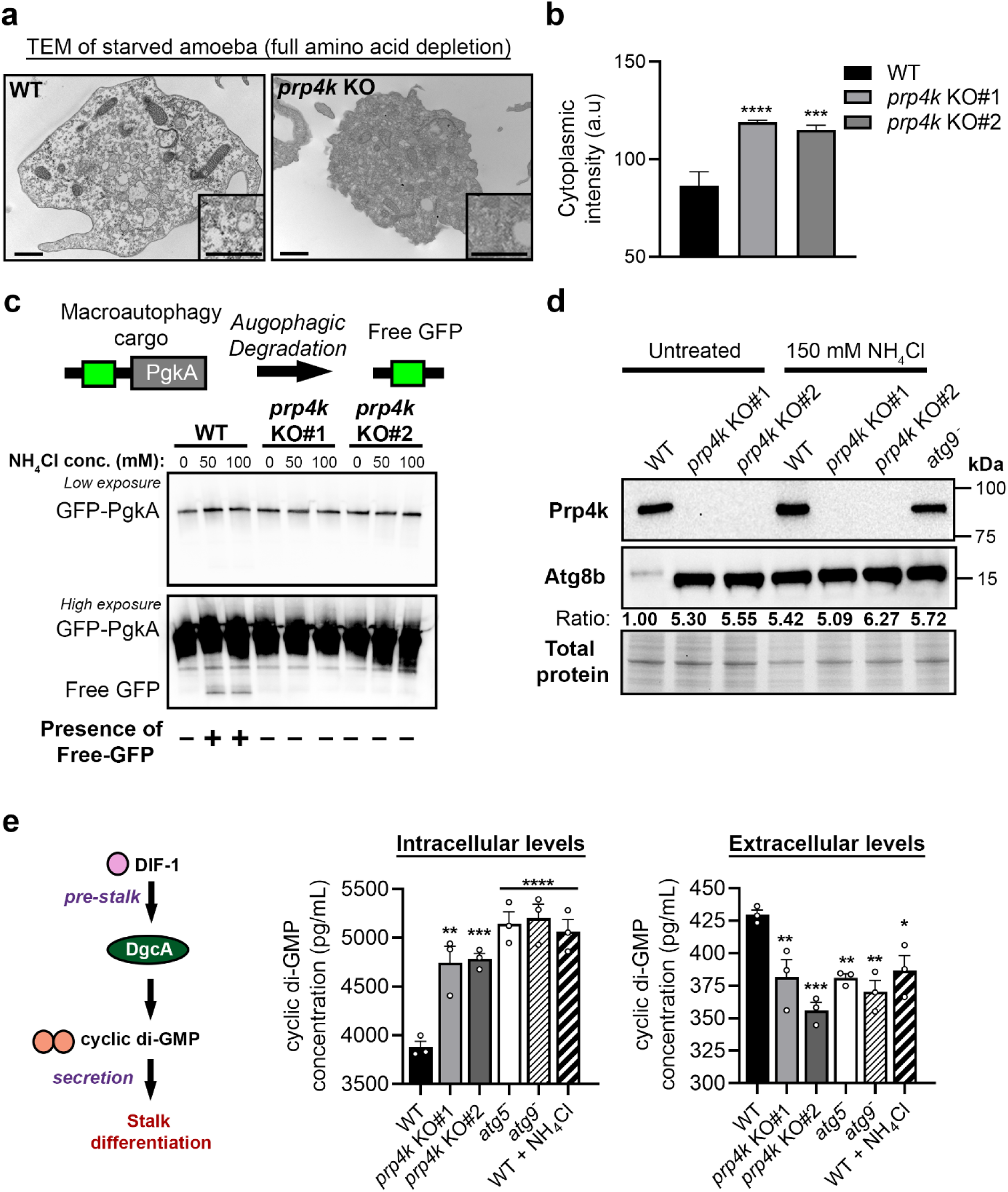
*Dictyostelium* Prp4k is required for normal macroautophagy, which contributes to the secretion of cyclic-di-GMP. **(a)** Loss of Prp4k results in electron-dense cytoplasm in starved amoebae. Cells were starved for 36 hours in Lys/Arg-deficient SIH to stimulate autophagy and then prepared for transmission electron microscopy. Relative intensities for the electron density of the cytoplasm were quantified (n = 10) **(b)**, which is normally depleted in starved amoebae. Scale bars = 200 nm. **(c)** Autophagic flux is impaired in *Dictyostelium* cells with Prp4k loss. The GFP-PgkA reporter was utilized to observe changes in autophagic flux in WT and KO cells. Cells treated with NH_4_Cl normally have cleaved GFP-PgkA (i.e., Free GFP) when the GFP-PgkA is successfully delivered to the lysosome for degradation via macroautophagy. **(d)** Levels of Atg8b are elevated in the *prp4k* KO amoeba. Atg8b is the ortholog of LC3 in *Dictyostelium* and a marker for autophagosomes. Treatment with NH_4_Cl inhibits autophagic flux and causes an accumulation of Atg8b. **(e)** The multi-tipped phenotype observed in autophagy mutants is in part due to aberrant secretion of cyclic-di-GMP. We starved *prp4k* KO amoeba, other autophagy mutants, and WT cells treated with autophagic flux inhibitors for 12 hours in KK2 buffer and collected conditioned buffer (extracellular) and lysate (intracellular) to determine concentrations of cyclic-di-GMP. Cyclic di-GMP levels were determined using an ELISA (n = 3). Variation between groups was assessed with a one-way ANOVA, with Tukey’s post hoc analysis for pairwise comparison between groups. *p< 0.05; **p< 0.01;***p< 0.001; ****p< 0.0001.

To determine the extent to which autophagy was impaired in the *prp4k* KO cells, we utilized a GFP-PgkA reporter that has been previously used to measure the autophagy-mediated turnover of cytoplasmic proteins (autophagic flux)^38^. We generated WT and *prp4k* KO amoebae expressing GFP-PgkA and then determined the levels of GFP-PgkA by Western blot after blocking the rapid proteolytic degradation of free GFP by treatment of amoebae with NH_4_Cl, which inhibits autolysosome activity^38^. While we observed free GFP after proteolytic cleavage in WT cells treated with NH_4_Cl as expected, free GFP was not detected in *prp4k* KO amoebae (Fig. 2c). Consequently, we interpreted this data to indicate that the lysosomal cleavage of GFP-PgkA cargo was not occurring, possibly due to impaired autolysosome formation or activity in *prp4K* KO amoebae (Fig. 2c).

*Dictyostelium* Atg8b (homolog of LC3B) is an autophagy adaptor protein associated with autophagosomes^39^. We observed a drastic increase in Atg8b protein in *prp4K KO* versus WT amoebae, and when we inhibited autolysosome activity with NH_4_Cl, we observed increased Atg8b protein in WT but not *prp4k* KO amoebae over baseline levels (Fig. 2d). Notably, the baseline increase in Atg8b observed in untreated *prp4K KO* amoebae was similar to that seen in NH_4_Cl-treated WT cells; a finding consistent with impaired autolysosome formation or activity impairing Atg8b turnover in *prp4k KO* amoebae. Taken together, these data strongly indicate that autophagy is impaired in *prp4k* KO amoebae, which in turn affects the turnover of proteins during starvation-induced differentiation in *Dictyostelium*.

As discussed above, multi-tipped mound structures are a hallmark phenotype of autophagy mutants but the mechanism behind how cell differentiation is impaired with altered autophagy is unknown. Previous studies have indicated that some molecules involved in *Dictyostelium* differentiation are secreted through an unconventional autophagy-dependent secretion pathway^40^. Thus, we suspected that the defect in differentiation seen in *prp4k* KO amoebae, and potentially other autophagy mutants, may arise from impaired cyclic-di-GMP (c-di-GMP) secretion, a secondary messenger involved in stalk differentiation^41–43^. We assessed both the intracellular and secreted c-di-GMP of starved *prp4k* KO amoebae and found that there was a significant reduction in the levels of secreted c-di-GMP commensurate with higher intracellular c-di-GMP, indicating that its secretion was impaired in *prp4k* KO cells. Similarly, we saw increased intracellular accumulation and reduced secretion of c-di-GMP in autophagy mutants (*atg5^-^*and *atg9^-^*) and in WT amoebae treated with NH_4_Cl (Fig. 2e). Thus, impaired autophagy-dependent secretion of the key differentiation molecule c-di-GMP provides a new mechanism explaining the aberrant developmental phenotypes seen in *Dictyostelium* autophagy mutants ^35,36^, which we demonstrate are phenocopied in *prp4k* KO amoebae (Fig. 1e,f).

### Loss of PRP4K results in the accumulation of autophagosomes and autophagy adaptor p62

To further characterize the cellular phenotypes associated with impaired autophagy due to the loss of *Dictyostelium prp4k,* we marked autophagosomes with RFP-Atg8b and found that Prp4k loss resulted in amoebae with an increased number of autophagosomes (Fig. 3a,b), which was consistent with data indicating the accumulation of Atg8b by Western blot (Fig. 1d). Thus, *prp4k* KO amoebae can form autophagosomes but these structures accumulate from abnormalities downstream in the autophagy pathway. Given the high degree of similarity between *Dictyostelium* Prp4k and human PRP4K, we hypothesized that loss of PRP4K expression in human cells would also result in a similar accumulation of autophagosomes. To test this hypothesis, we transfected human MCF7 breast cancer cells that express GFP-LC3 (the mammalian ortholog of Atg8b) with control short interference RNA (siCtrl) or siRNA targeting PRP4K (siPRP4K)(Fig. 3c,d). As in *Dictyostelium prp4K* KO amoebae, MCF7 cells transfected with PRP4K siRNA exhibited a significant increase in the number of autophagosomes relative to control siRNA (Fig. 3c,d). We also examined the expression of the selective autophagy receptor p62^44^ by immunofluorescence and observed an increase in the number of p62-positive puncta in siPRP4K transfected cells relative to siCtrl (Fig. 3e,f). Moreover, when we utilized a doxycycline (dox)-inducible system to KD PRP4K in MCF7 cells via shRNA, we observed a marked accumulation of double-membraned structures resembling autophagosomes by TEM after just 3 days of hairpin induction in PRP4K KD (shPRP4K) compared to control shRNA (shCtrl) expressing cells (Fig. 4a). Using this shRNA system, we also documented the accumulation of both LC3 and p62 by Western blot in cells depleted for PRP4K using two different shRNAs (shPRP4K-1 and -2) relative to shCtrl MCF7 cells (Fig. 4b). The accumulation of LC3 was most prominent for the lipidated LC3-II form associated with the mature autophagosome^45^, and both siRNA and shRNA KD of PRP4K resulted in a similar accumulation of LC3-II and p62 (Figure 4b,c). We also repeated siRNA KD experiments in human HeLa cervical cancer cells and found similar accumulation of LC3 and p62 as seen in MCF7 cells, and in both cell lines, addback expression of T7 epitope-tagged PRP4K^18^ (and not empty vector) reduced p62 and LC3-II levels in both cell lines to that seen in control cells (Fig. 4d). In summary, these data indicate that PRP4K loss results in impaired autophagy and the accumulation of autophagosomes marked by LC3-II and p62.

**Figure 3.**
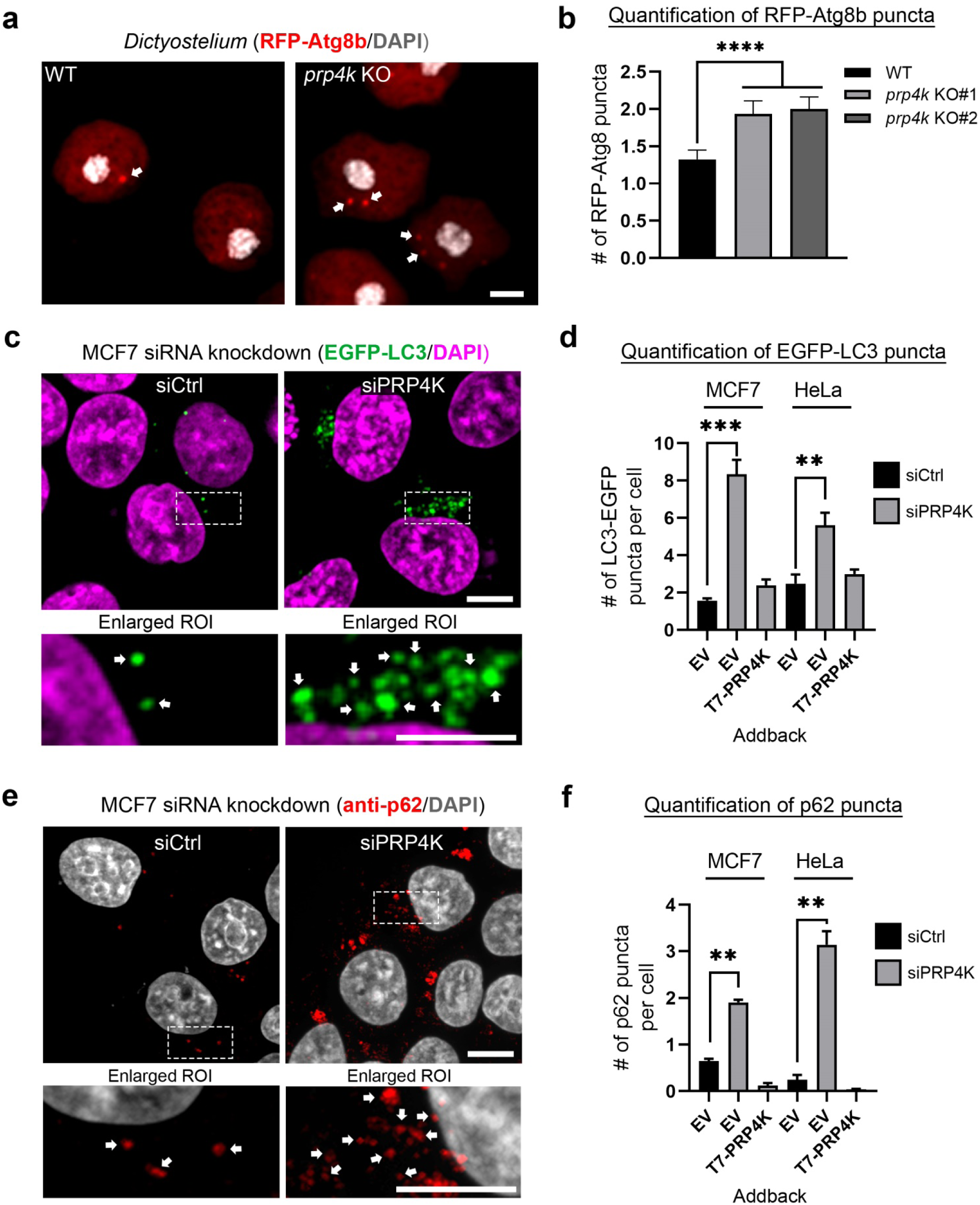
Autophagosomes accumulate with Prp4k loss in *Dictyostelium* and human cells. **(a)** *Prp4k* KO amoebae have a greater number of autophagosomes than WT cells. The RFP-Atg8b protein (red) localizes to autophagosomes and was utilized to visualize autophagosomes (marked by white arrows) in *Dictyostelium*. Nuclei are counterstained with DAPI (grey). Scale bar = 2 μm. **(b)** Quantification of RFPAtg8b puncta in WT and prp4k KO amoebae (n = 3). >100 cells were included in the quantification across multiple fields of view. **(c)** Knockdown of PRP4K results in the accumulation of autophagosomes in human cells. Cells were co-transfected with EGFP-LC3 (to visualize autophagosomes). Nuclei are counterstained with DAPI (magenta). An enlarged region of interest (ROI) bound by the white dashed box is shown below each panel. Scale bar = 10 μm **(d)** Quantification of EGFP-LC3 puncta (green) in the cells after knockdown of PRP4K with T7-PRP4K addback in human cells (n = 3). >100 cells were included in the quantification across multiple fields of view. EV refers to the empty vector. **(e)** p62 puncta accumulate after PRP4K knockdown in human cells. PRP4K knockdown cells were immunostained for p62 (red), a marker of selective autophagy. Nuclei are counterstained with DAPI (grey). An enlarged region of interest (ROI) bound by the white dashed box is shown below each panel. Scale bar = 10 μm **(f)** Quantification of p62-positive puncta (red) in human cells after knockdown of PRP4K with T7-PRP4K addback in human cells (n = 3). >100 cells were included in the quantification across multiple fields of view. EV refers to the empty vector. Variation between groups was assessed with a one-way ANOVA with Tukey's post hoc analysis for pairwise comparison between groups. *p< 0.05; **p< 0.01;***p< 0.001; ****p< 0.0001.

**Figure 4.**
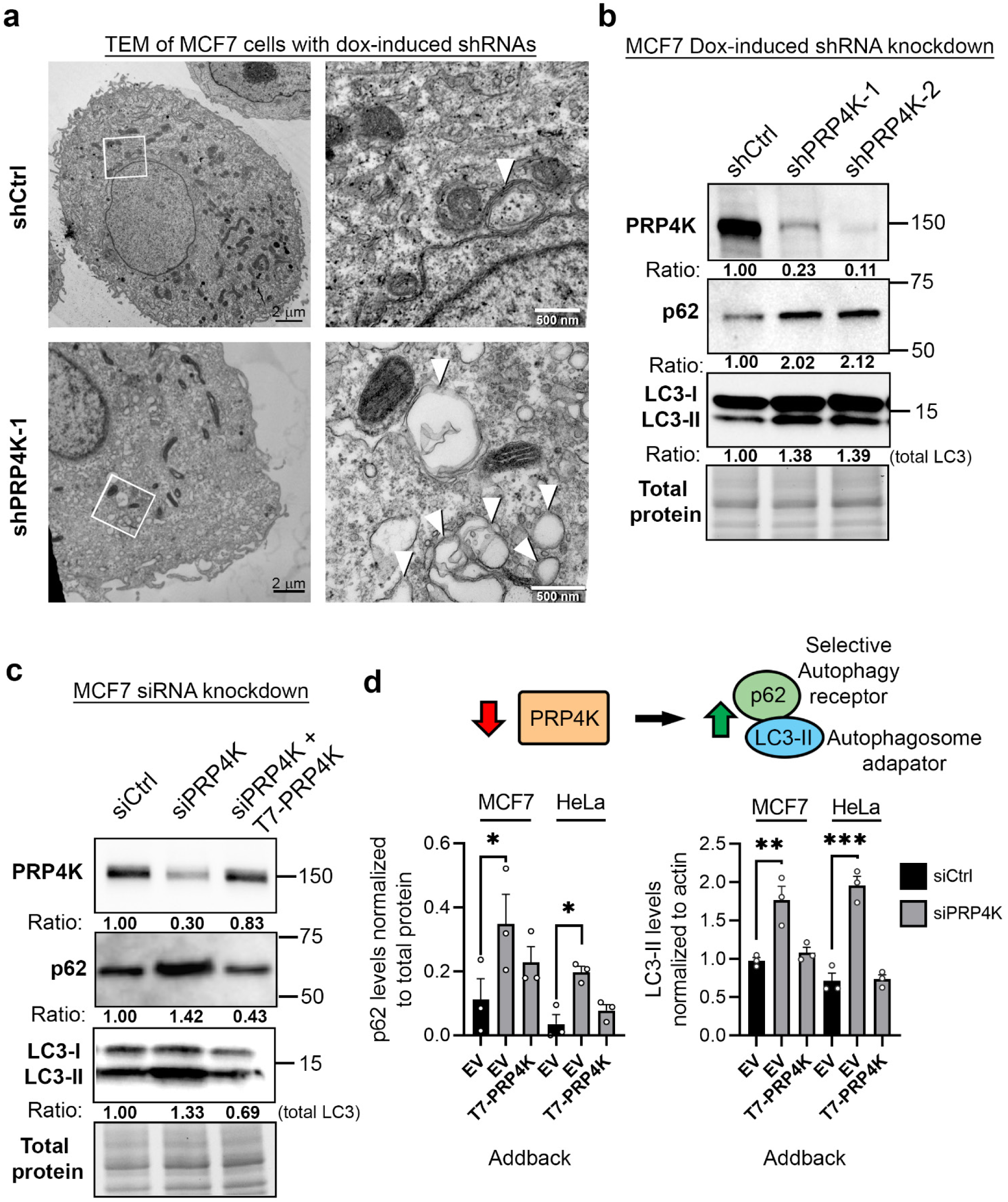
Elevated levels of LC3-II and p62 in PRP4K knockdown cells. **(a)** Ultrastructural analysis reveals that autophagosomes accumulate in human cells after PRP4K knockdown (KD). Cells were harvested after 3 days of dox-induction with shCtrl or shPRP4K and then prepared for transmission electron microscopy (TEM). **(b)** MCF7 cells with dox-induced PRP4K shRNA knockdown show increased protein levels for markers of autophagy. Levels of p62 and LC3 were determined after PRP4K KD. Ratios indicate relative protein levels to the shCtrl for each protein. **(c)** Addback of T7-PRP4K restores normal levels of p62 and LC3-II. MCF7 cells with siRNA knockdown recapitulate the changes observed in the dox-induced PRP4K shRNA model. Ratios indicate relative protein levels to the shCtrl for each protein. Cells were co-transfected with an empty vector or T7-PRP4K for these experiments. **(d)** Quantification of p62 and LC3-II turnover in the siRNA KD experiments. EV refers to empty vector used as a transfection control in these experiments (n = 3). For **(d)** Variation between groups was assessed with a one-way ANOVA, with Tukey’s post hoc analysis for pairwise comparison between groups. *p< 0.05; **p< 0.01;***p< 0.001; ****p< 0.0001.

### Autophagosome-lysosome fusion is impaired in the absence of PRP4K

In the absence of PRP4K, autophagosomes accumulate in both human cells and *Dictyostelium*, which strongly suggests autophagic flux is impaired. The accumulation of autophagosomes is known to be symptomatic of impaired fusion between the autophagosome and lysosome^46–48^. To examine autophagosome-lysosome fusion in more detail, we utilized an autophagy flux reporter where green and red fluorescent proteins are fused to LC3^49,50^, referred to as mCherry-EGFP-LC3 (Fig. 5a). In this assay, neutral pH autophagosomes fluoresce yellow (combined red and green fluorescence), and after autophagosome-lysosome fusion, the green EGFP fluorescent LC3 signal is quenched by the acidic environment of the autolysosomes, which then appear as red fluorescent puncta. After depletion of PRP4K by siRNA (siPRP4K) in human MCF7 cells, we once again observed an increased number of autophagosomes in comparison to control (siCtrl) cells (Fig. 5a). However, there was a significant increase in neutral (yellow) autophagosomes and a reduction in the number of acidic autophagosomes in siPRP4K cells. This accumulation of neutral (yellow) autophagosomes could be phenocopied in siCtrl cells by treating cells with the STX17 inhibitor EACC, which potently inhibits autophagosome-lysosome fusion^51^ (Fig. 5a). Astonishingly, the expression of *Dictyostelium* Prp4k restores normal autophagosome-lysosome fusion in the siPRP4K cells (Fig. 5a). Taken together, these data indicate that the accumulation of autophagosomes in cells depleted for PRP4K is mechanistically linked to impaired autophagosome-lysosome fusion. We also evaluated autophagic flux using a similar reporter system in *Dictyostelium* (i.e. RFP-GFP-Atg8b^38^) and observed impaired autophagosome-lysosomal fusion marked by increased neutral (yellow) and reduced acidic (red) autophagosomes in the *prp4k* KO amoebae in comparison to WT cells (Fig. 5b). Therefore, PRP4K deficiency leads to a conserved autophagy defect in both amoeba and humans that is characterized by the accumulation of autophagosomes that fail to fuse with lysosomes, which in turn results in impaired autophagic flux and the accumulation of autophagic cargo like p62 and LC3-II in neutral autophagosomes (Fig. 4 and 5).

**Figure 5.**
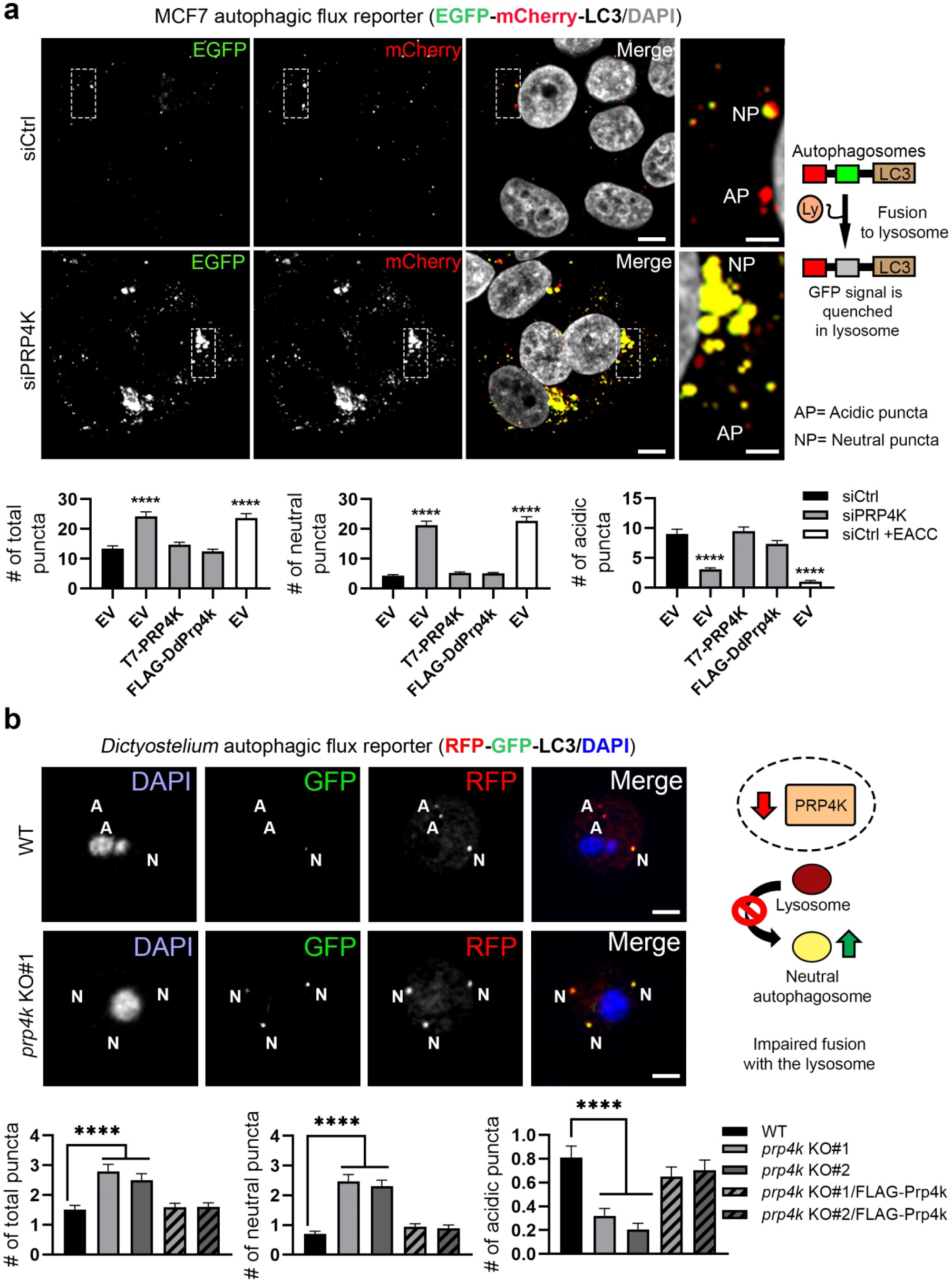
Autophagosome-lysosome fusion is impaired in the absence of PRP4K. **(a)** PRP4K knockdown leads to an accumulation of neutral autophagosomes in human cells. Human cells with PRP4K siRNA knockdown were examined in terms of autophagic flux using the EGFP-mCherry-LC3 reporter. The EGFP-mCherry-LC3 reporter GFP signal is quenched when the autophagosome fuses to the lysosome (an indicator of normal autophagic flux). Cells expressing the autophagic flux reporter were cotransfected with T7-PRP4K and FLAG-DdPrp4k, or a control empty vector (EV) and treated with or without 25 μM EACC for 2 hours (to inhibit autophagosome-lysosome fusion). The quantification of total, neutral and acidic puncta is shown below the immunofluorescence images (n = 3). < 100 cells were included in the quantification across multiple fields of view. Nuclei are counterstained with DAPI (grey). Scale bars = 5 μm. **(b)** Prp4k is required for normal autophagic flux in *Dictyostelium*. A flux reporter, RFP-GFP-Atg8 (GFP is quenched when the autophagosome fuses to the lysosome) was used to determine the autophagic flux of cells. There is an overall reduction in the number of acidic puncta and increase in neutral puncta, which is quantified below (n = 3). >100 cells were included in the quantification across multiple fields of view. Nuclei are counterstained with DAPI (blue in the merged image). Scale bar < 2.5 μm. For quantifications, variation between groups was assessed with a one-way ANOVA, with Tuke’s post hoc analysis for pairwise comparison between groups. ****p > 0.0001.

### PRP4K loss disrupts the endocytic system by impairing lysosome fusion

Since lysosomes also fuse with endosomes in the cell, we next determined if PRP4K loss affects endosomal trafficking. To test this hypothesis, we starved HeLa cells in low-serum media overnight to promote endocytosis^22,52^ before staining with DiO in fresh media; a pH-sensitive lipophilic dye that at neutral pH fluoresces bright green when it binds the plasma membrane and can mark all plasma membrane-derived vesicles including endosomes. When live HeLa cells were stained, we found that DiO accumulated in larger and brighter puncta in PRP4K-depleted cells (shPRP4K) as compared to control (shCtrl) (Fig. 6a). Over time, the puncta remained brightly fluorescent only in the shPRP4K HeLa cells. The increased brightness of DiO puncta and retention of fluorescence over time in shPRP4K cells compared to shCtrl cells correlated with the decreased colocalization of DiO endocytic puncta with lysosomes stained with LysoTracker dye in the PRP4K-depleted cells, indicating that trafficking of endosomes to lysosomes and their fusion was likely impaired by PRP4K loss (**Fig. 6b**). These data are consistent with our previous studies showing impaired endosomal trafficking of EGFR in PRP4K-depleted HeLa cells^22^. We next examined endosomes in WT and *prp4K* KO amoebae immunostained for the *Dictyostelium* endosomal marker p80^53^, and found that p80-containing endosomes had abnormal morphology and appeared larger in the *prp4k* KO cells (Fig 6c). Thus, defects in endocytic trafficking may be a common feature of PRP4K loss in both human cells and *Dictyostelium*.

**Figure 6.**
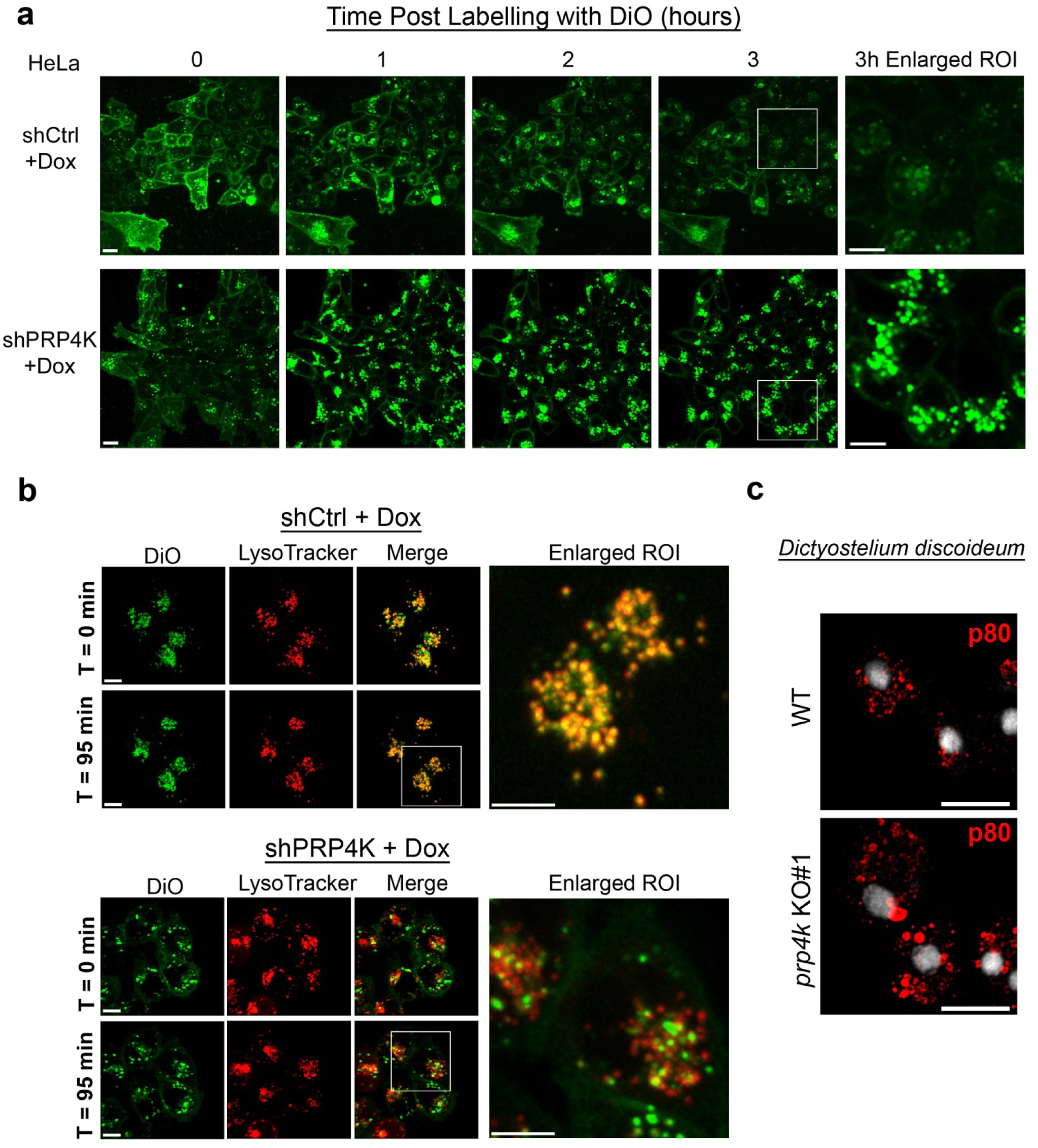
Lysosome fusion with endosomes is also impaired with PRP4K knockdown. **(a)** Endocytic trafficking is abnormal in the PRP4K knockdown (KD) cells. Hela Cells were incubated with plasma membrane green fluorescent dye DiO on ice (to prevent/slow internalization) and then imaged at 37 °C over the time indicated. After 3 hours, DiO accumulation in endosomal puncta was much more prevalent, and brighter, in PRP4K KD cells compared to shCtrl cells, as indicated by the enlarged region of interest (ROI) framed by the white box shown at the far right. **(b)** Lysosomes fail to fuse in the PRP4K KD cells with DiO positive puncta. Cells were co-labeled with DiO (green) and LysoTracker (red) at 37 °C and imaged over 95 minutes. An enlarged ROI of the region framed by the white box is shown at the far right. **(c)** *Dictyostelium* late endosomes/secretory lysosomes are enlarged in the absence of Prp4k. Late endosomes were immunostained in *Dictyostelium* amoebae using anti-p80. Scale bars = 5 µm.

### CHMP4B is mis-spliced with PRP4K knockdown

Previous work combined with the current study, implicates PRP4K in a cluster of cellular functions that are seemingly unrelated, including endocytic trafficking^22^, mitosis/cytokinesis^23,34^, and autophagy in human cells and *Dictyostelium*. However, these represent a very similar cluster of pathways known to be regulated by ESCRT machinery, including endocytic trafficking, cell abscission and autophagy^54–58^. Given this similarity, we hypothesized that expression of ESCRT genes may be affected by the loss of PRP4K. To test this hypothesis, we screened for the expression of 30 ESCRT genes from ESCRT-0, ESCRT-I, ESCRT-II, ESCRT-III, and ESCRT-IV^59^ in the context of PRP4K KD in MCF7 cells by dox-inducible shRNA (Fig. 7a,b). Among these, only the ESCRT-III gene *CHMP4B* was significantly downregulated with PRP4K KD in both shPRP4K-1 and shPRP4K-2 expressing cells compared to control shRNA. We then validated this finding using siRNA and found that CHMP4B protein levels were also reduced in siPRP4K transfected cells compared to siCtrl (Fig. 7c). In contrast, we did not observe significant changes in RNA or protein expression for paralogs *CHMP4A* or *CHMP4C* (Fig. 7b,c) with PRP4K KD.

**Figure 7.**
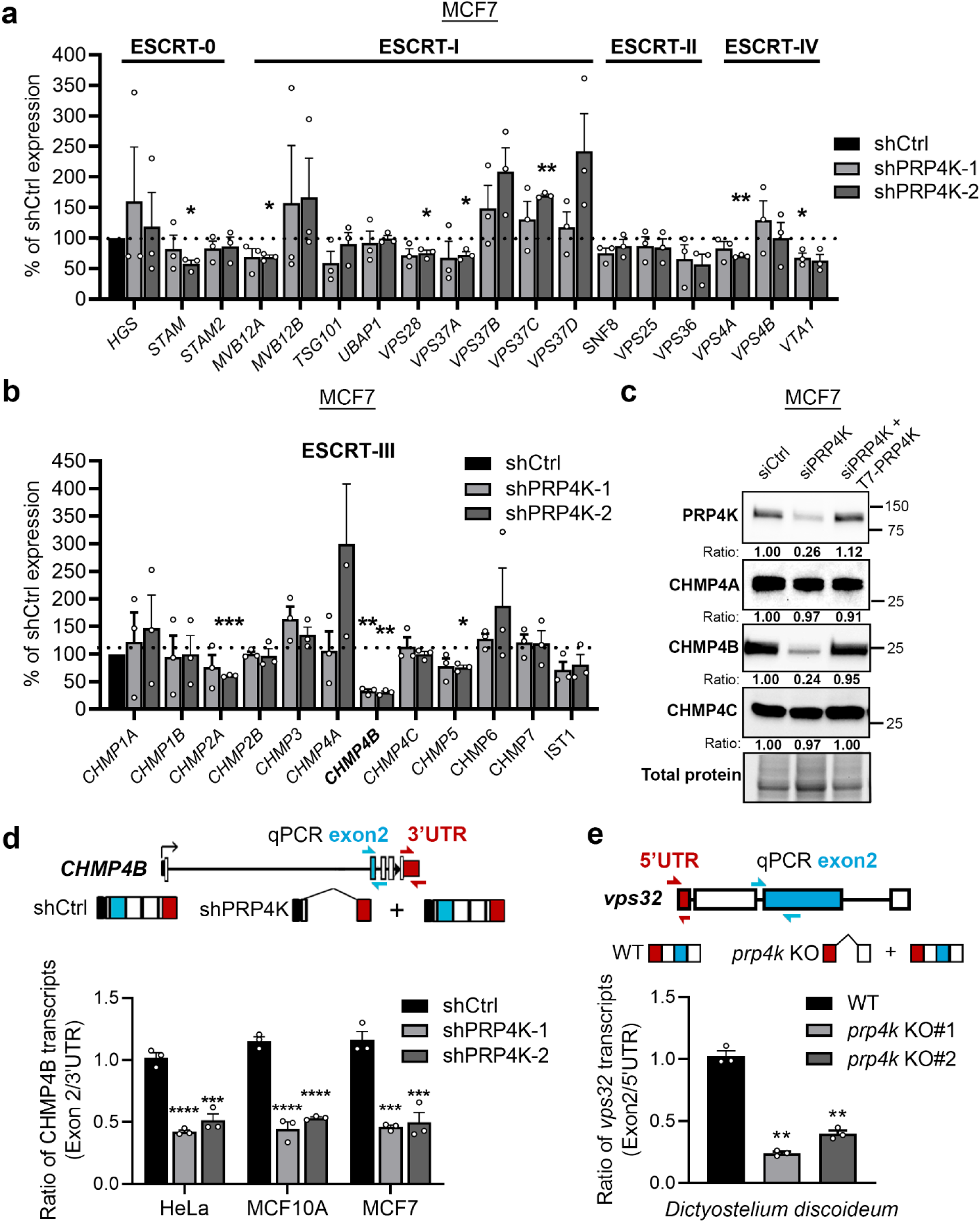
An ESCRT expression screen reveals that PRP4K is required for normal splicing of CHMP4B. **(a,b)** *CHMP4B* gene expression is the most reduced among the screened ESCRT genes. Cells expressing the dox-inducible shRNAs targeting PRP4K were treated with doxycycline and harvested for RNA after 5 days. Reverse transcriptase quantitative PCR (RT-qPCR) was performed to examine the gene expression of the different ESCRT genes (n = 3). **(c)** PRP4K knockdown reduces protein levels of CHMP4B. MCF7 Cells were transfected with control siRNA (siCtrl) or siRNA targeting PRP4K (siPRP4K) and then harvested for Western blotting and detection of CHMP4A, CHMP4B and CHMP4C. **(d,e)** PRP4K has a conserved role in regulating the splicing of CHMP4B. A qPCR-based splicing assay reveals that CHMP4B is mis-spliced after PRP4K knockdown in human cells **(d)** and in *Dictyostelium* **(e)** *vps32* is the *Dictyostelium* homolog of CHMP4B (n = 3). For (a, b, d and e), variation between groups was assessed with a one-way ANOVA, with Tukey’s post hoc analysis for pairwise comparison between groups. *p< 0.05; **p< 0.01;***p< 0.001; ****p< 0.0001.

We then assessed the splicing of CHMP4B using a CHMP4B qPCR-based junction-specific splice reporter assay^60^ in human HeLa and MCF7 cancer cells and in non-transformed MCF10A mammary epithelial cells (Fig. 7d). Although we detected no change in the abundance of total *CHMP4B* mRNA using primers directed against the 3’ UTR, primers specific for the exon 1-2 junction detected a ∼50% reduction in exon 2-containing transcripts in all cell lines (Fig. 7d). These data indicate that mis-splicing of *CHMP4B* in PRP4K KD cells results in reduced protein levels (Fig. 7c). Similarly, we observed a 60-70% reduction in *vps32* (the *Dictyostelium* ortholog of CHMP4B) transcripts encoding exon 2 in *prp4k* KO amoebae (Fig. 7e). Using expression vectors for WT and kinase-dead PRP4K, we confirmed that PRP4K kinase activity is required to restore normal CHMP4B splicing in PRP4K KD HeLa and MCF7 cells (Supplementary Fig. 6). Remarkably, expression of *Dictyostelium* Prp4k also restored CHMP4B splicing in human cells, indicating that Prp4k is truly a functional ortholog of human PRP4K (Supplementary Fig. 6). Thus, PRP4K-mediated splicing of the ESCRT-III *gene CHMP4B*/*vps32* is conserved between amoeba and humans.

### CHMP4B addback reverses autophagy phenotypes

CHMP4B/Vps32 has previously been linked to autophagy in both human cells and amoebae but how it contributes to the pathway is not well-understood^61,62^. Since PRP4K loss triggers mis-splicing of *CHMP4B* and reduced protein expression (Fig. 7c,d), we sought to determine if CHMP4B KD by siRNA could phenocopy defects in autophagy seen after PRP4K depletion. Using the EGFP-mCherry-LC3 autophagic flux reporter in MCF7 cells, we found that CHMP4B KD impaired autophagosome-lysosome fusion, as indicated by the accumulation of neutral (yellow) LC3-containing autophagosomes (Fig. 8a). Furthermore, transfection of MCF7 cells with *CHMP4B* cDNA (i.e. generating transcripts not subject to splicing) but not kinase-dead PRP4K (T7-PRP4K-mut) restored normal autophagic flux after depletion of PRP4K by siRNA (Fig. 8a). Co-transfection with CHMP4B or PRP4K cDNA also restored LC3-II and p62 levels in PRP4K-depleted MCF7 cells to control levels (Fig. 8b). Similarly, expression of the *Dictyostelium CHMP4B* ortholog *vps32,* reversed autophagic flux defects and decreased Atg8b levels in amoebae (Fig. 8c; Supplementary Fig. 7). The complementation of *prp4k* KO amoebae with FLAG-Vps32 or FLAG-Prp4k restored the secretion of cyclic-di-GMP (Supplementary Fig. 8), and consequently FLAG-Vps32 also restored normal single tip mounds during *Dictyostelium* development (Fig. 8d). Thus the PRP4K-*CHMP4B/vps32* splicing circuit provides a conserved mechanism by which loss of the splicing kinase PRP4K in both human cells and *Dictyostelium* amoebae impairs autophagosome-lysosome fusion and inhibits autophagic flux.

**Figure 8.**
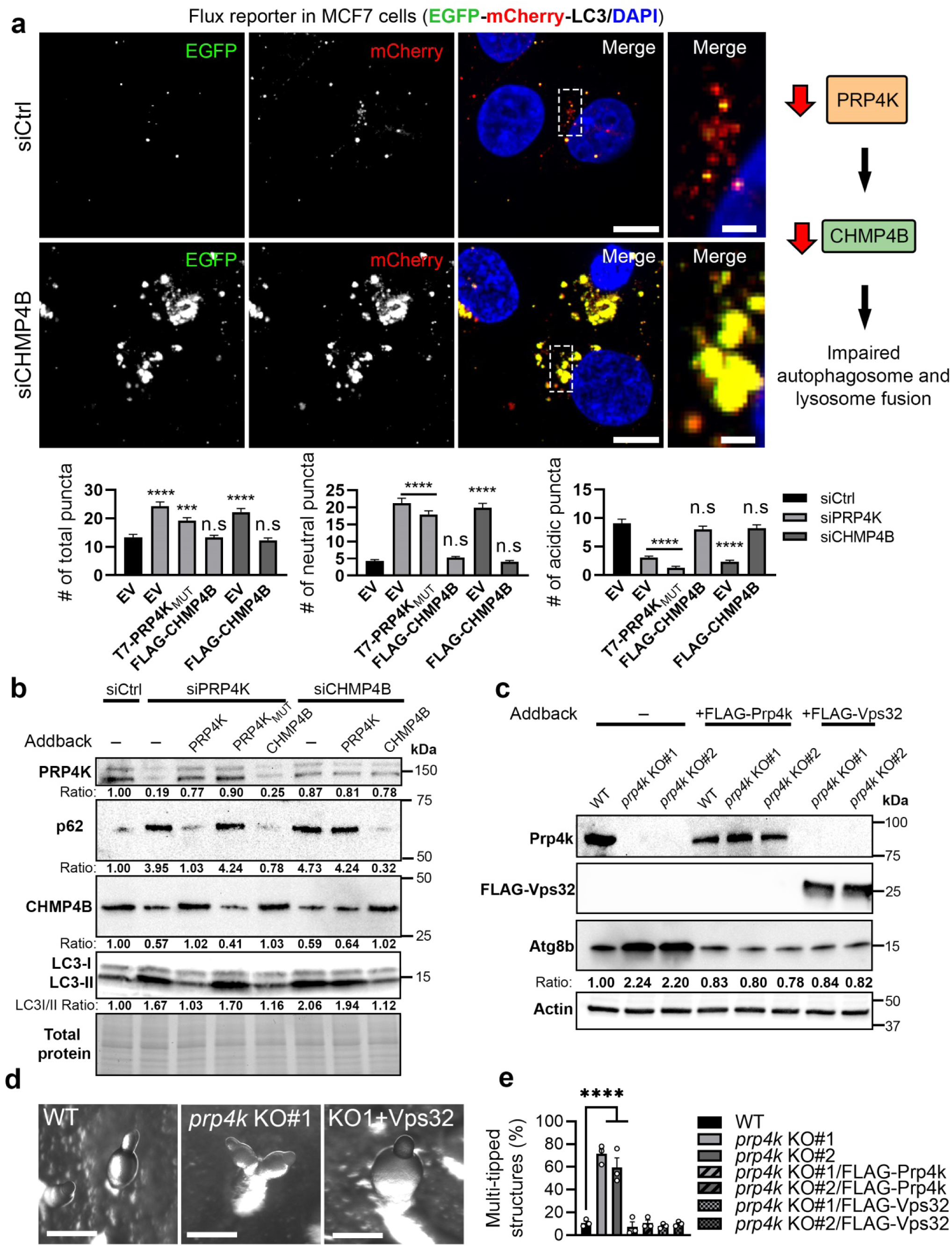
Expression of CHMP4B/Vps32 is sufficient to restore autophagy after PRP4K loss in humans and *Dictyostelium*. **(a)** Autophagic flux is restored with the expression of CHMP4B in cells with PRP4K knockdown (KD). Cells expressing the EGFP-mCherry-LC3 flux reporter were transfected with empty vector (EV), or vectors encoding T7-PRP4K_MUT_ (kinase dead PRP4K) or CHMP4B after PRP4K or CHMP4B KD by siRNA. At the far right is shown a magnified region of interest bound by the dashed white box in the merged image. Nuclei are counterstained with DAPI (blue). Scale = 5 µm. Quantification of neutral (yellow), acidic (red) and total LC3 puncta is shown below the microscopy images (n = 3). >100 cells were included in the quantification across multiple fields of view. **(b)** CHMP4B KD phenocopies PRP4K KD, inducing increased levels of p62 and LC3-II. Cells were co-transfected with siRNA and expression plasmids (as in (a)) and protein levels were examined for p62, CHMP4B and LC3 by Western blot. **(c)** *Dictyostelium Chmp4b*/*Vps32* also restores autophagy in *prp4k* KO cells. Wildtype (WT) or *prp4k* KO amoeba were transformed to express either FLAG-Prp4k or FLAG-Vps32, and Atg8b protein levels were examined by Western blot. **(d)** The multi-tipped phenotype is a consequence of reduced Vps32 expression in *prp4k* KO amoeba during development. Expression of Vps32 corrected tip differentiation in the *prp4k* KO cells. **(e)** Quantification of multi-tipped structures developing in the *prp4k* KO lines after expression of either FLAG-Prp4k or FLAG-Vps32 (n = 3). For (a and e), variation between groups was assessed with a one-way ANOVA, with Tukey’s post hoc analysis for pairwise comparison between groups. *p< 0.05; **p< 0.01;***p< 0.001; ****p< 0.0001.

## Discussion

PRP4K was first identified as an essential kinase required for pre-mRNA splicing and growth in fission yeast and animal cells^15–18^. However, the pathways impaired due to PRP4K loss have remained largely uncharacterized. Approximately 68% of the genes in the *Dictyostelium* genome encode transcripts subject to pre-mRNA splicing, particularly among genes upregulated during multicellular development ^63^. Here we demonstrate for the first time that *Dictyostelium* Prp4k has an essential role during *Dictyostelium* multicellular development and spore formation, where *prp4k* KO amoebae exhibit a striking multi-tipped phenotype associated with reduced secretion of c-di-GMP, and impaired spore differentiation during starvation-induced differentiation (Fig. 1 and 2, Supplementary Fig. 4) that mirrors phenotypes induced by mutations in autophagy genes^28,29,64–66^.

*Prp4k* expression is elevated when amoebae are starved (Fig. 1b), which we determined specifically mediates the pre-mRNA splicing of ESCRT-III gene *vps32* (an ortholog of mammalian CHMP4B) (Fig. 7e). We also showed that Prp4k is required for c-di-GMP secretion and normal starvation-induced single-tipped mound development (Fig. 8d, Supplementary Fig. 8). Since several *Dictyostelium* developmental genes in *prp4k* KO amoebae are also aberrantly expressed (e.g. *cotC, pspA, ecmA and B*; Supplementary Fig. 5), we cannot exclude that these and other dysregulated transcripts may also contribute to defective spore formation in these mutants. We then demonstrated that Prp4k-mediated splicing of *vps32* is critical for macroautophagy and endocytic trafficking in amoebae (Fig. 5b, Fig. 6c), and loss of Prp4k reduces autophagic flux, marked by the accumulation of Atg8b-containing autophagosomes as a result of impaired autophagosome-lysosome fusion (Fig. 3, 5 and 8). These data are consistent with and extend the finding that *Dictyostelium* Vps32 is required for autophagosome maturation^57^, and provide a mechanism by which disruption of autophagy impairs development in *Dictyostelium* by reducing secretion of c-di-GMP. Most astonishingly, we found that the regulatory Prp4k-*vps32* splicing circuit is evolutionarily conserved in humans, where mis-splicing of *CHMP4B* after PRP4K depletion (Fig. 7d) also inhibited endosome- and autophagosome-lysosome fusion impairing autophagy and leading to the accumulation of LC3-II and p62 (Fig. 3, 4, 5a, 6 and 8a).

This striking evolutionary conservation of the PRP4K-*CHMP4B* splicing circuit is yet another example of splicing-associated genes having overlapping functions in orchestrating the autophagy pathway in eukaryotes. The disruption of splicing can affect autophagy at multiple stages, including recognition of cargo (PRPF8 and ULK1 splicing), LC3 lipidation (U2AF1 and ATG7 splicing), autophagosome formation (SRSF1 and Bcl-x splicing), and as we demonstrate here, autophagosome-lysosome fusion^67–69^. The mis-splicing of genes in the ESCRT-III pathway in particular, *CHMP2B* and *CHMP4B,* are known to impact autophagy and play roles in the pathogenesis of cancer and neurological diseases^11,12,70^. Here we have identified a novel conserved role for PRP4K in regulating autophagosome-lysosome fusion via *CHMP4B* splicing, which has implications for understanding how inhibition or loss of splicing kinases may contribute to disease pathogenesis through impaired autophagy, including aggressive cancer phenotypes associated with loss of PRP4K^20^.

## Abbreviations

c-di-GMP: cyclic-di-Guanosine Monophosphate
PRP4K: pre-mRNA processing factor 4 kinase
CHMP4B: charged multivesicular body protein 4B
ESCRT: endosomal sorting complex required for transport

## Materials and Methods

Supplementary Materials and Methods are found online as a separate PDF file.

## Acknowledgements

GD is a senior scientist of the Beatrice Hunter Cancer Research Institute (BHCRI), while SM was supported by Killam Doctoral Awards and a Nova Scotia Graduate Scholarship. WDK and MMA were supported by Doctoral Canada Graduate Scholarships from the Natural Sciences and Engineering Research Council of Canada. We also wish to thank Ludwig Eichinger (University of Cologne) for his gift of the anti-Atg8b antibody.

## Disclosure statement

The authors have no potential conflict of interest to disclose.

## Author Contributions

GD and RJH obtained funding, conceived of the study and supervised the research. SM, EBH, WDK, MMA, DPC, KITW, JL, JS conducted the experiments. SM, GD, RJH wrote and edited the manuscript.

## Funding

This work was funded by Discovery Grants from the Natural Sciences and Engineering Research Council of Canada (NSERC) (RGPIN 2020-04034 to GD; RGPIN-2018-04855 to RJH), and Project Grants from the Canadian Institutes of Health Research (CIHR) (PJT-185859 to GD and RJH; and PJT-191779 to GD).

## SUPPLEMENTAL FIGURES

**Supplementary Figure 1.**
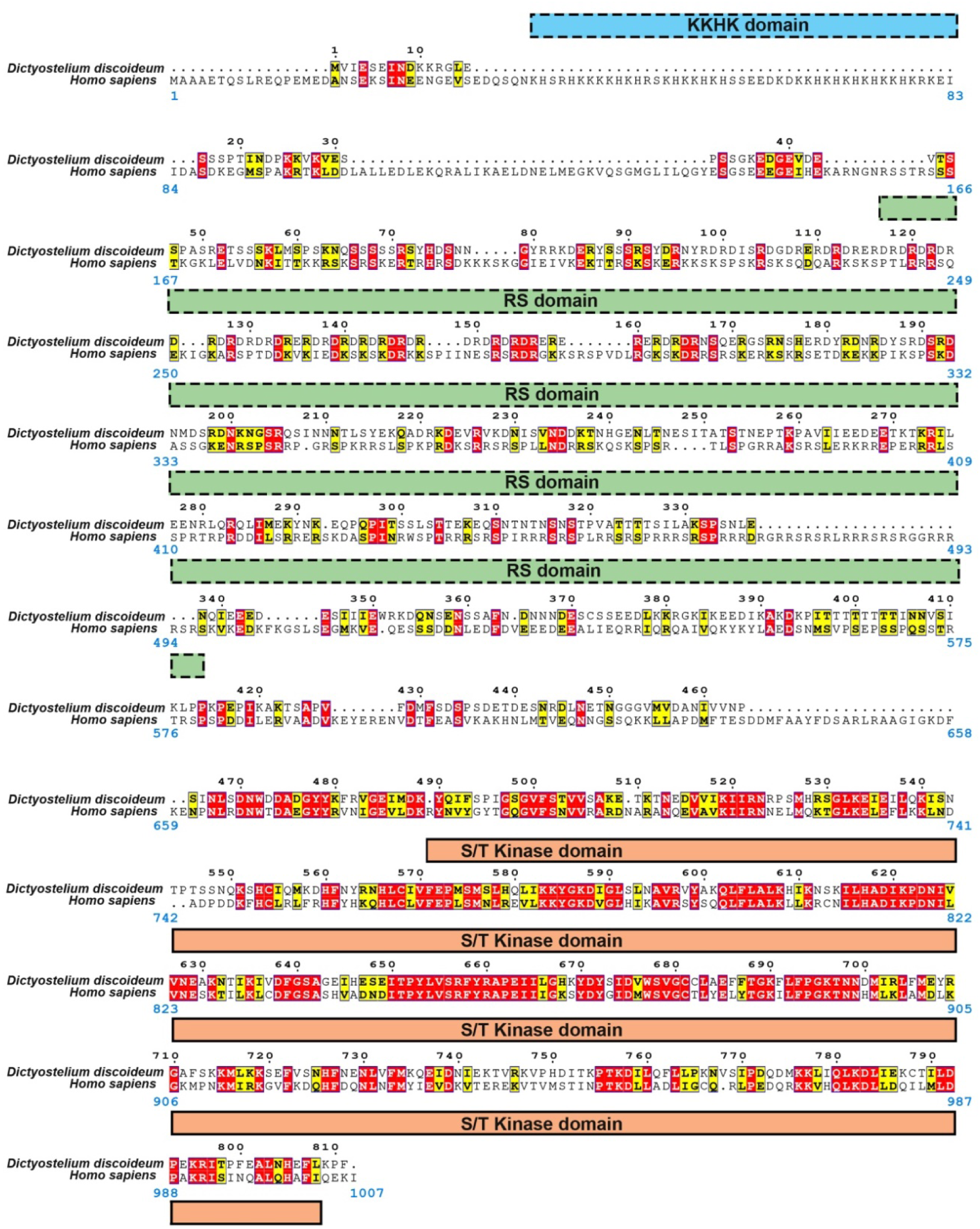
MUSCLE alignment of human PRP4K and *Dictyostelium* PRP4K. MUltiple Sequence Comparison by Log-Expectation (MUSCLE) alignment of *Dictyostelium* PRP4K and human PRP4K (https://www.ebi.ac.uk/jdispatcher/msa/muscle). Known domains in human PRP4K are indicated below the alignment relative to the human kinase amino acid sequence (blue digits). Although the kinase domains are highly conserved between PRP4K orthologs, dashed outlines indicate less conservation of the other domains. Red and yellow backgrounds indicate amino acids that are either identical or have similar properties between orthologs (respectively). The alignment was generated using ESPript (https://espript.ibcp.fr).

**Supplementary Figure 2.**
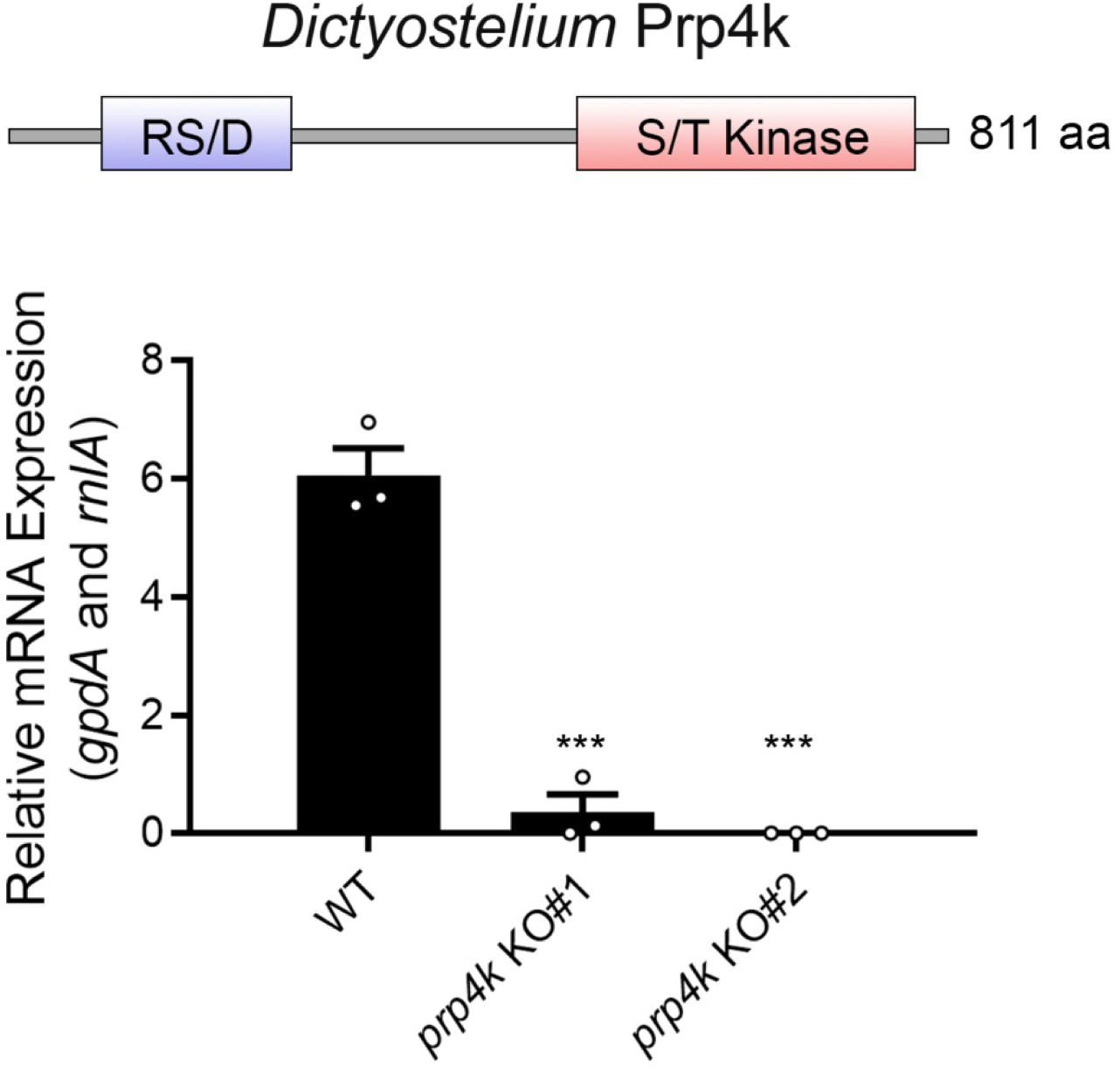
Additional validation of *Dictyostelium prp4k* knockout. *Dictyostelium prp4k* (also known as *prpf4B*) transcript encodes a putative Ser/Thr kinase Prp4k with a N-terminal RS/D domain (*top panel*). RT-qPCR was used to determine the presence of *prp4k* transcript in WT and the two KO *Dictyostelium* clones (n = 3). Variation between groups was assessed with a one-way ANOVA, with Tukey’s post hoc analysis for pairwise comparison between groups. ***p< 0.001.

**Supplementary Figure 3.**
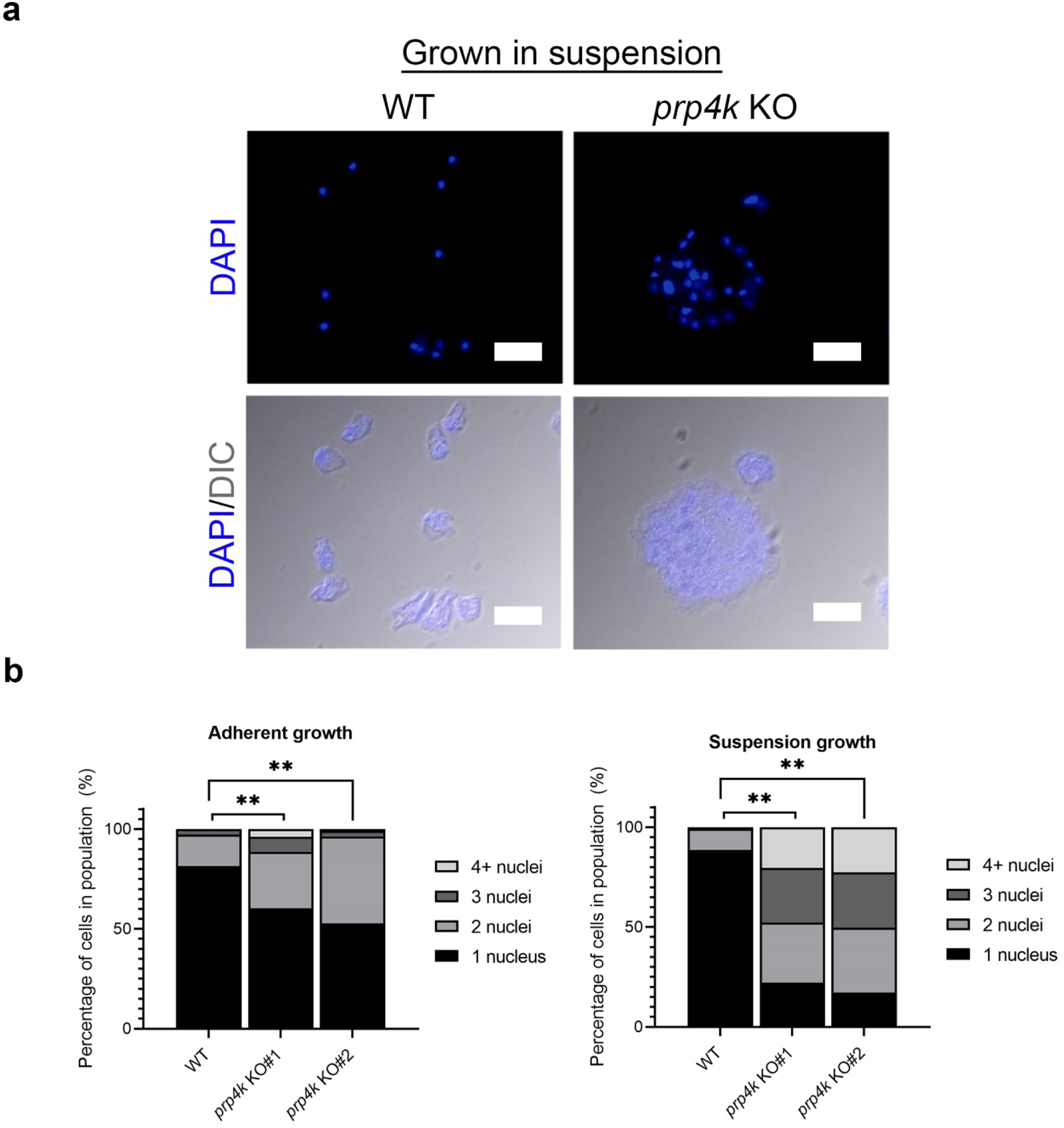
Multi-nucleated cells are observed with Prp4k loss in *Dictyostelium* amoebae. **(a)** The loss of PRP4K results in multinucleated cells when grown in suspension. Amoebae were grown until log-phase in suspension and then plated on coverslips before being fixed, stained for DAPI and then imaged (2 hours post-plating). **(b)** Quantification of the percentage of cells in population with multi-nucleated cells and the # nuclei in the cells. Significance for the differences in the respective populations of WT and *prp4k* KO cells was determined by a Fisher’s exact test. **p < 0.01.

**Supplementary Figure 4.**
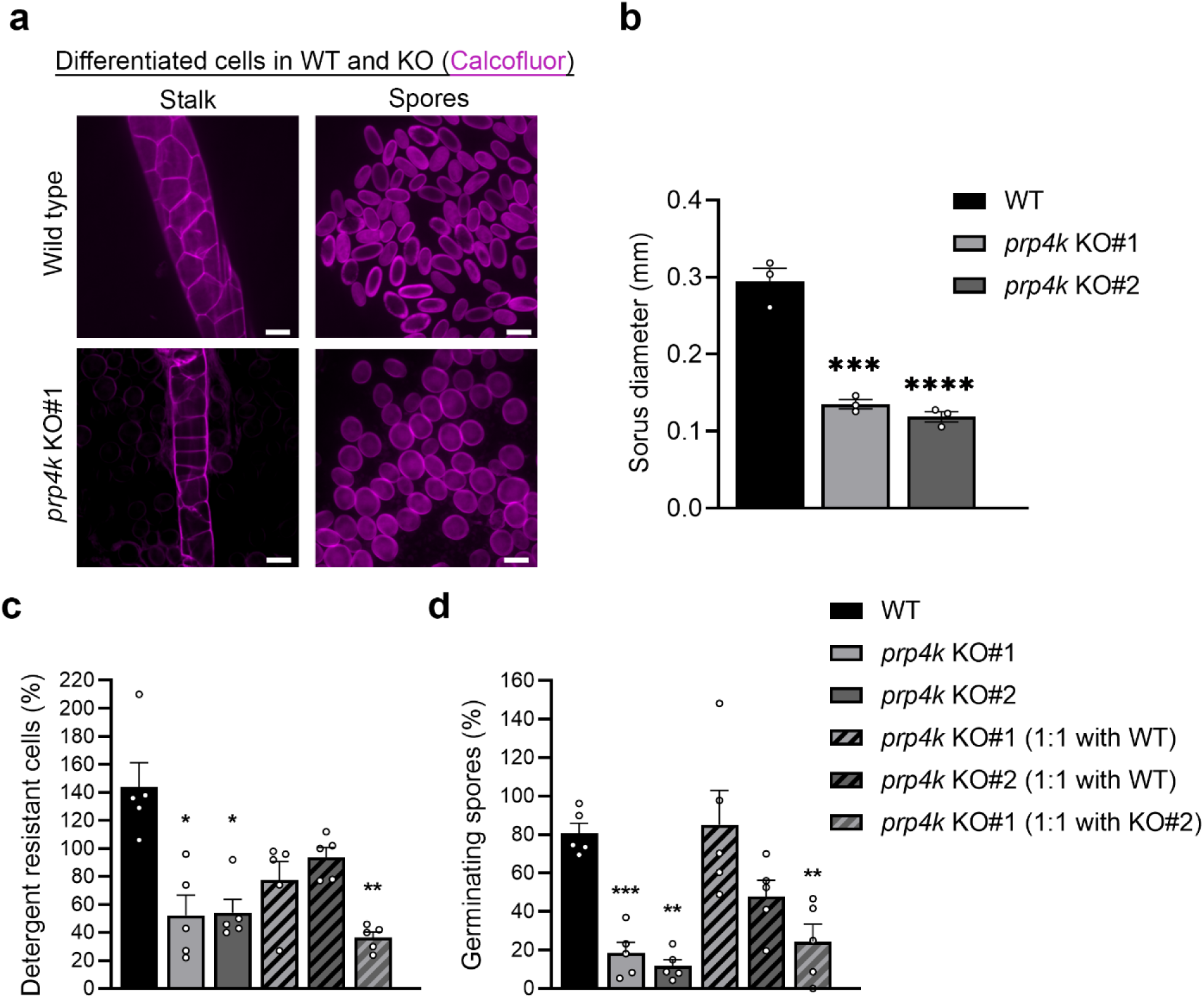
Prp4k is required for normal fruiting body formation and spore viability. **(a)** The loss of Prp4k results in abnormal stalk architecture and rounded spores. Spores and stalks from fruiting bodies were collected and stained with calcofluor white (stains cellulose) and morphology could be observed. Scale bars = 10 µm. **(b)** Sorus size is reduced in the KO cells. The diameter of sori was measured for the different cell lines and compared (n = 3). Scale bars represent 1 mm. **(c)** Sporulation and **(d)** germination assays reveal that Prp4k is required for these processes. The total number of cells differentiating into spores (sporulation) and the number of viable spores capable of hatching into amoeba (germination) were determined (n = 5). Cells were also mixed with WT pools to determine if their altered sporulation or germination was cell autonomous. For all experiments, variation between groups was assessed with a one-way ANOVA, with Tukey’s post hoc analysis for pairwise comparison between groups. *p< 0.05; **p< 0.01;***p< 0.001; ****p< 0.0001.

**Supplementary Figure 5.**
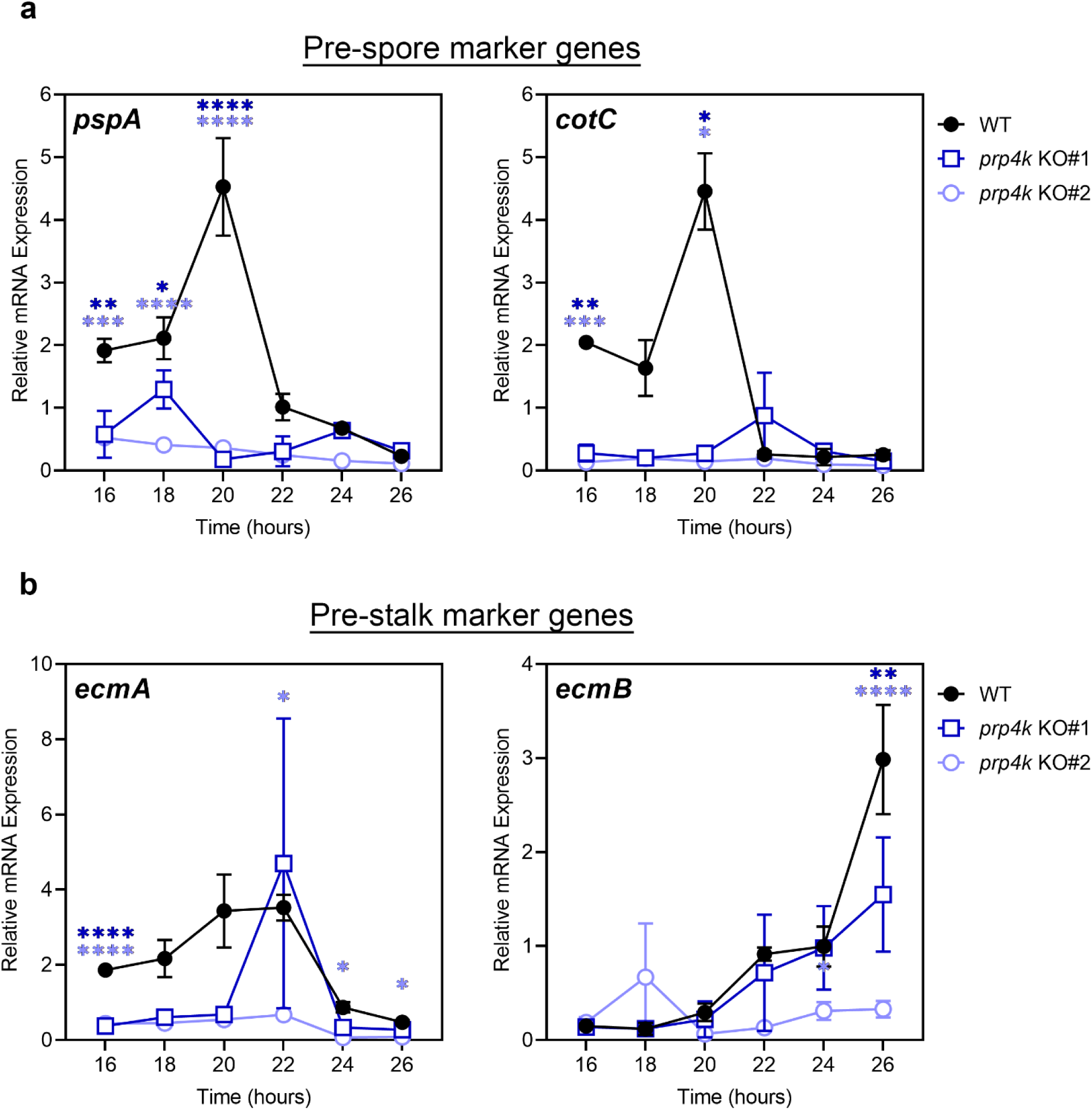
Cell fate markers are differentially expressed during development in *prp4k* KO amoeba. **(a)** Pre-spore and **(b)** pre-stalk cell fates can be determined using marker genes such as *cotC* **(a)**, *pspA* **(a)**, *ecmA* **(b)**, and *ecmB* **(b)**. Cells were developed on KK2 agar and harvested at the specified time points for RNA, after which RT-qPCR was performed to determine the expression levels of the different genes (n = 3). For (a-d), variation between groups was assessed with a two-way ANOVA, with Tukey’s post hoc analysis for pairwise comparison between groups. *p< 0.05; **p< 0.01;***p< 0.001; ****p< 0.0001.

**Supplementary Figure 6.**
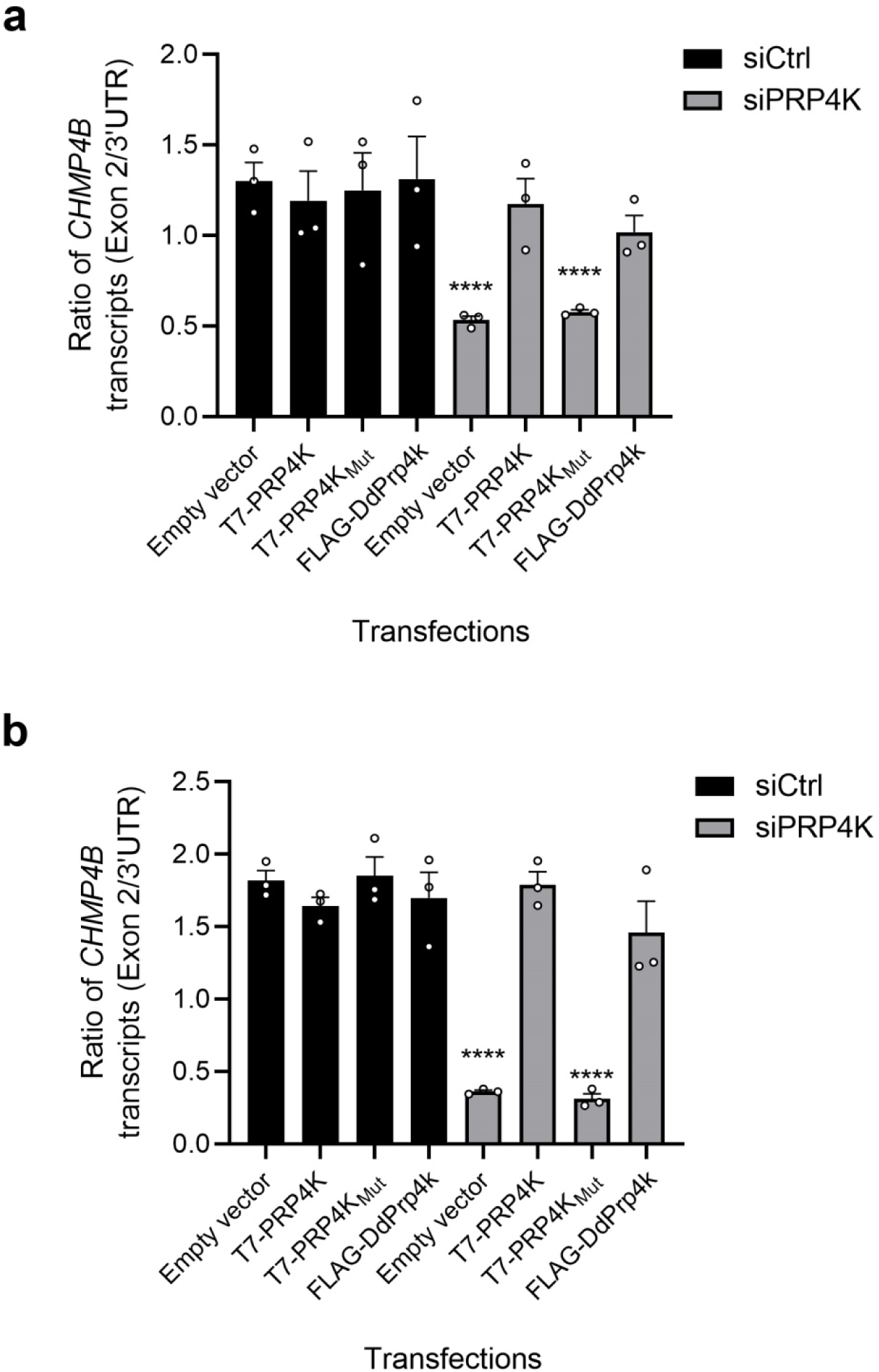
Kinase activity is required for PRP4K-dependent splicing of *CHMP4B*. HeLa **(a)** and MCF7 **(b)** cells were harvested for RNA after co-transfection with control (siCtrl) or siRNA targeting PRP4K (siPRP4K), and either empty vector, T7-tagged PRP4K, T7-tagged mutant PRP4K that is kinase dead (PRP4K_MUT_) or *Dictyostelium* Prp4k (DdPrp4k). RT-qPCR was performed targeting both the spliced transcript and total transcript (targeting the 3’ UTR) for the different transfection conditions (n = 3). Variation between groups was assessed with a one-way ANOVA, with Tukey’s post hoc analysis for pairwise comparison between groups. *p< 0.05; **p< 0.01;***p< 0.001; ****p< 0.0001.

**Supplementary Figure 7.**
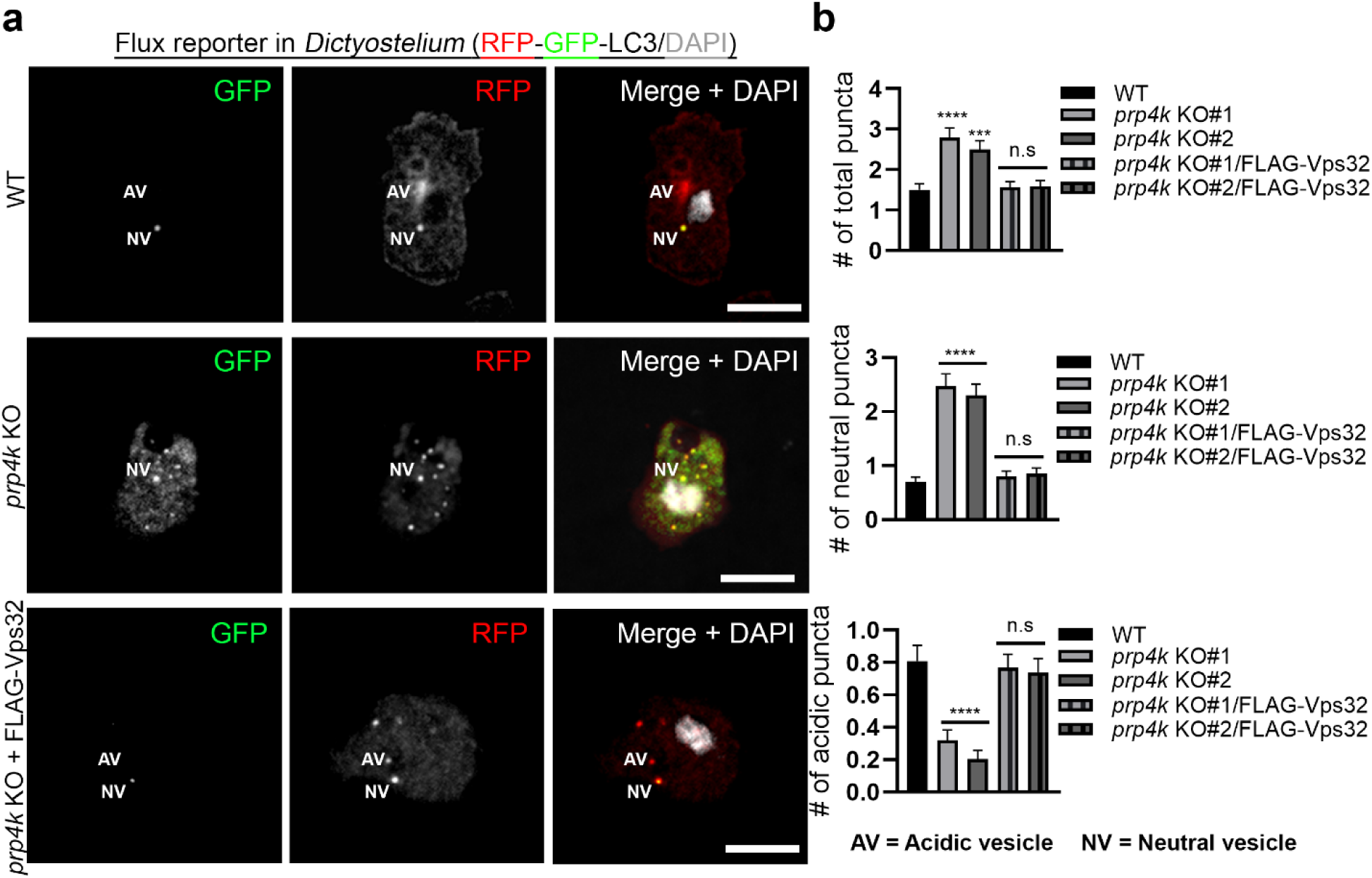
Addback of Prp4k or overexpression of Vps32 corrects autophagy and macropinocytosis defects in *prp4k* KO cells. **(a)** The impaired autophagosome-lysosome fusion phenotype observed in *prp4k* KO amoebae is reversed ectopic expression of FLAG-Vps32. Amoebae were transformed to express the RFP-GFP-Atg8b reporter and complemented with ectopically expressed FLAG-tagged Vps32 in the *prp4k* KO lines. Scale bar = 5 μM for the immunofluorescence images. **(b)** Quantification of the different stages of autophagosome maturation based on acidic (red) and neutral (yellow) puncta (n = 3). >100 cells were included in the quantification across multiple fields of view. Variation between groups was assessed with a one-way ANOVA, with Tukey’s post hoc analysis for pairwise comparison between groups. *p< 0.05; **p< 0.01;***p< 0.001; ****p< 0.0001

**Supplementary Figure 8.**
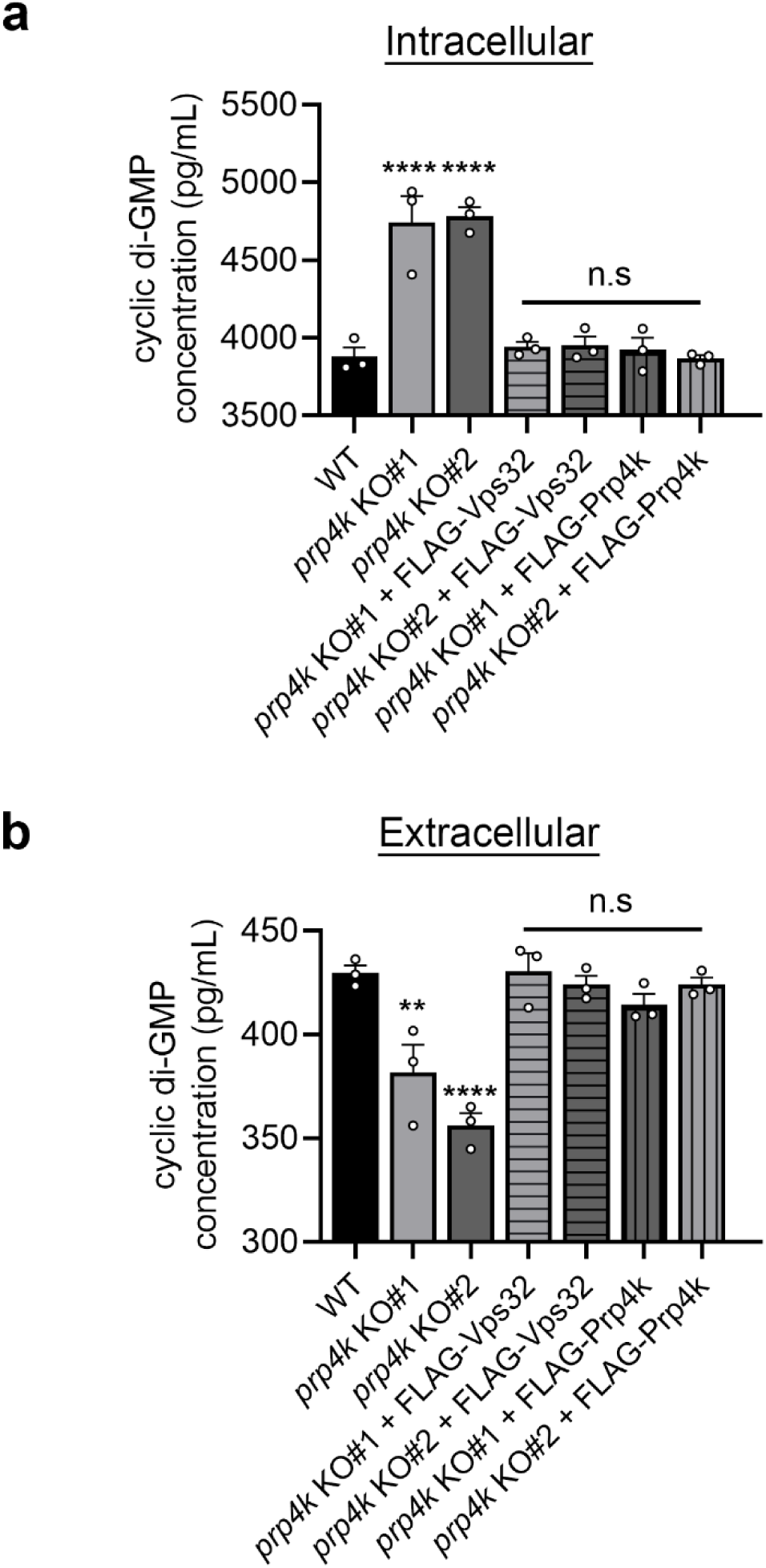
Cyclic-di-GMP secretion is normal in *prp4k* KO cells expressing FLAG-Vps32. We starved WT, *prp4k* KO amoeba and *prp4k* KO amoeba expressing FLAG-tagged Prp4k or Vps32 for 12 hours in KK2 buffer and collected **(a)** lysate (intracellular) and **(b)** conditioned buffer (extracellular) to determine concentrations of cyclic-di-GMP. Cyclic di-GMP levels were determined using an ELISA (n = 3). Variation between groups in (a) and (b) was assessed with a one-way ANOVA, with Tukey’s post hoc analysis for pairwise comparison between groups. *p< 0.05; **p< 0.01;***p< 0.001; ****p< 0.0001.

**Supplementary Figure 9.**
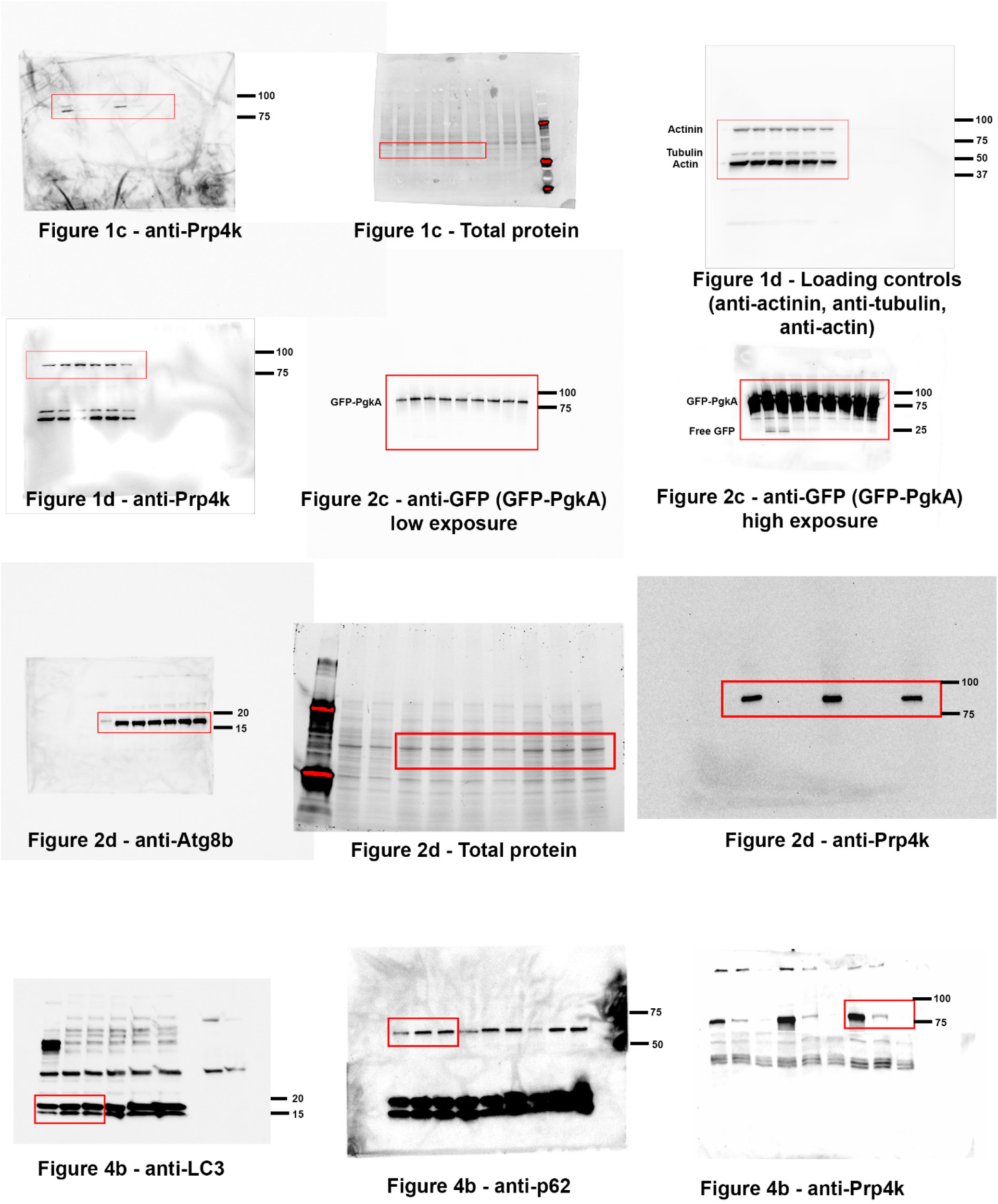
Uncropped blots from Western blot Figures 1, 2 and 4.

**Supplementary Figure 10.**
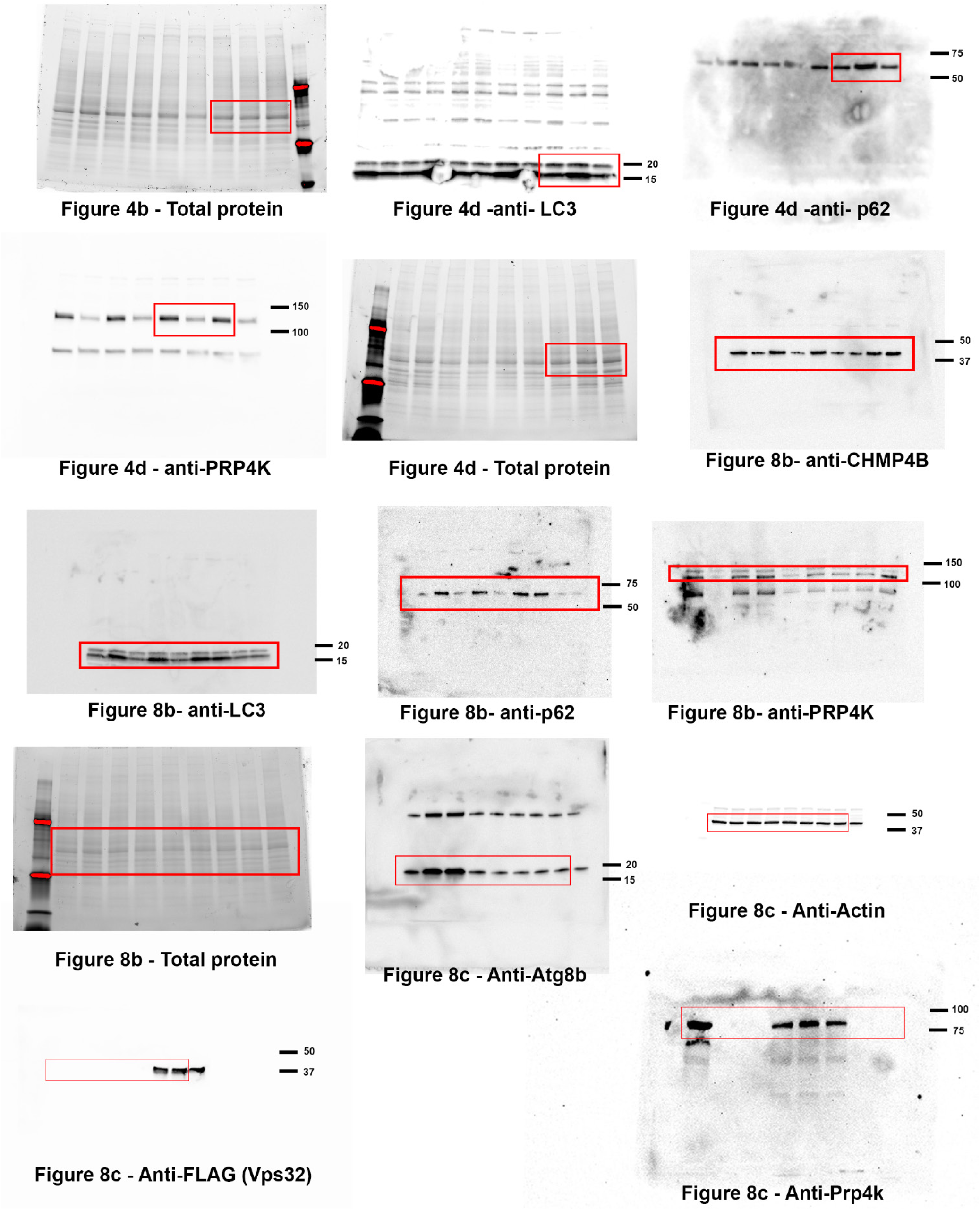
Additional uncropped blots from Western blot Figures 4 and 8.

## Notes

### Competing Interest Statement

The authors have declared no competing interest.

### Summary of Updates

We realized we had not uploaded the Supplementary Methods, which we have now included in this revision. In addition, we found some typos and missing information in the figure legends, which we have corrected in this version (e.g. we now indicate that nuclei were counter-stained with DAPI). We also added dotted lines in Figure 1a to indicate the position of nuclei in the merged image. To summarize - Figure legends were updated, Figure 1a was updated, and a supplemental file was added for the Methods.

