## Supplementary Materials and Methods for "The evolutionarily conserved PRP4K-*CHMP4B/vps32* splicing circuit regulates autophagy"

*Dictyostelium Cell Lines, Media, and Buffers*

*Dictyostelium* cells were grown and maintained on SM/2 agar with *Klebsiella aerogenes* at 22°C as previously described[1]. Liquid cultures were supplemented with ampicillin (100 µg/ml) and streptomycin sulfate (300 µg/ml) (Bioshop Canada Incorporated, Burlington, ON, Canada). The cells used for experiments were grown axenically in nutrient-rich medium (HL5) at 22°C under adherent conditions on plastic Petri dishes. HL5, minimal medium (FM), FM lacking amino acids (FM-aa), and low-fluorescence HL5 were purchased from Formedium (Hunstanton, Norfolk, United Kingdom). AX3 cells, referred to as WT cells, were the parental cell line for the *prp4k* KO mutant cells. For assessing autophagy in *Dictyostelium*, WT and *prp4k KO* cells were transformed to express pA15/GFP-Atg8a (marks autophagosomes), pA15/GFP-PgkA (autophagy reporter) and pA15/RFP-GFP-Atg8a (autophagic flux reporter) [2] using vectors purchased from the Dicty Stock Center[6]. Cell lines were generated using a transformation protocol described previously[3]. Briefly, cells were washed with ice-cold H50 buffer, concentrated to 1 x $10^7$ cells/mL and 100 µL of cells were mixed with 1-2 µg of plasmid DNA before being electroporated using the Gene Pulser Xcell™ Electroporation System with 0.1 cm cuvettes. *atg5*$^{-}$ and *atg9*$^{-}$ cells were purchased from the Dicty Stock Center [4-6].

*Human Cell Lines, Medias and Buffers*

HeLa cell lines were cultured in Dulbecco's Modified Eagle's Medium (DMEM; Sigma) supplemented with 10% fetal bovine serum (FBS, Wisent) and 1% penicillin/streptomycin (P/S) at 37°C with 5% $CO_2$. MCF7 human breast cancer cells were cultured in Eagle's Minimal Essential Medium (MEM; Sigma) using T-75 flasks, supplemented with 10% FBS, 1% penicillin/streptomycin (P/S), 10 µg/mL human recombinant insulin (Sigma 19278), and 100X

GlutaMAX™ (ThermoFisher Scientific, catalog# 35050061) at 37°C with 5% $CO_2$. Cell lines stably expressing shRNAs were cultured as mentioned above but using 1 μg/mL puromycin (ThermoFisher, A1113802). To inhibit autophagosome-lysosome fusion, cells were treated for 2 hours with 25 μM EACC (catalog# HY-129111, MedChemExpress).

*Generation of the CRISPR KO* in *Dictyostelium*

To generate a *prpf4k* knockout (KO) cell line, we used the pTM1285 all-in-one sgRNA CRISPR/Cas9 expression vector, as previously described[7]. The pTM1285 vector was kindly provided by the University of Tsukuba through the National Bio Resource Project (NBRP) of the MEXT Japan. Candidate single guide-RNA (sgRNA) sequences for the *prpf4b* gene were designed using CRISPOR and ordered containing overhangs specific to BpiI (Concordet and Haeussler, 2018). Two sets of sgRNAs were designed, Set 1 (Sense: 5' - AGCAAGTGGGTTCATTAGTAGATG – 3'; Antisense: 5' – AAACCATCTACTAATGAACCCACT – 3') and Set 2 (Sense: 5' - AGCACTCTGTTGTAGATAAGGAAG – 3'
Antisense: 5' – AAACCTTCCTTATCTACAACAGAG – 3'). Single sgRNAs were cloned into the pTM1285 vector using BpiI according to manufacturer's instructions (New England Biolabs, Whitby, ON, Canada). Ligation of the sgRNA was confirmed through PCR with the sense oligo sgRNA from the 2 selected sgRNAs (sequences below) and the tracrRV primer 5'-AAGCTTAAAA AAAGCACCGACTCGGTGCC-3'. Sanger sequencing was conducted prior to transformation into *D. discoideum* with the tracrRV primer (5'-AAGCTTAAAAAAAGCACCGACTCGGTGCC-3').

Validated pTM1285-sgRNA constructs were transformed into AX3 cells via electroporation using established *D. discoideum* protocols[3]. Cells in the mid-log phase of growth were pelleted at 500 g for 5 minutes and washed twice in ice-cold H50 buffer (20 mM HEPES, 50 mM KCl, 10 mM NaCl, 1 mM Mg2SO4, 5 mM NaHCO3, 1 mM Na2HPO4). The cells were resuspended in a total volume of 100 µL ice-cold H50 buffer containing 6 µg of validated pTM1285-sgRNA construct. A 0.1 cm cuvette was pre-chilled prior to use, and the cell suspension was added to the cuvette, incubated on ice for 5 minutes and then electroporated using a MicroPulser electroporator (Bio-Rad Laboratories Canada, Mississauga, ON, Canada) with the *Dictyostelium* settings according to the Bio-Rad instruction manual (1 kV, 2 pulses, 1 ms time constant). The cuvette was placed immediately back into ice and let sit for 15 minutes after adding a solution of CaCl2 and MgCl2 to a final concentration of 1 mM for each respective compound. The cells were then plated in 100 mm x15 mm Petri dishes with 10 mL HL5 containing ampicillin (100 µg/mL) streptomycin (300 µg/mL) and left to sit at room temperature for 8 hours. G418 was added to each Petri dish at a final concentration of 20 µg/mL, and the plate was left for four days. Following the four-day incubation, the remaining cells were counted, washed, serially diluted, and plated (100 cells) onto SM/2 agar plates with *Klebsiella aerogenes*.

Individual clones (plaques) were isolated from the agar plates and transferred to 12-well plates containing HL5 with only ampicillin and streptomycin (between four to seven days following plating on SM). The clones transferred to the 12-well plates were slow to grow and took over a week to reach 70% confluency in axenic culture in the 12-well dish. Therefore, we did not scale growth up to large petri dishes in axenic culture. The cells were washed from the 12-well plate in 100 µL KK2 buffer, and 2 µL of the cell suspension was collected, washed 2x in KK2 and then lysed for DNA extraction using 48 µL LyseB buffer (10 mM Tris, pH 8.3, 50 mM

KCl, 2.5 mM MgCl2, 0.45% NP40, and 0.45% Tween) and 2 μL Proteinase K (Charette and Cosson, 2004). The remaining cells were transferred to SM/2 agar plates with KA to grow and then store for future use at -80C. To detect indels generated from CRISPR/Cas9, an 876 bp region spanning the sgRNA cut site was amplified via PCR and ran on an agarose gel. PCR amplicons from clones with visible indels were sent for Sanger sequencing, and we selected 2 clones to work with - one with a 7-bp insertion (+GTGTTCA at nucleotide position 758 of the coding region of prpf4b and the other a 4-bp deletion (-CCTC at nucleotide position 899 of the coding region of prpf4b) that resulted in a frameshift and early truncation of the gene product.

*Dictyostelium multicellular development*

Multicellular development of *Dictyostelium* cells was examined as previously described [35]. Briefly, cells grown in HL5 and heat killed bacteria were harvested from Petri dishes and washed two times with ice-cold 1x KK2 phosphate buffer (2.2 g/L $KH_2PO_4$, 0.7 g/L $K_2HPO_4$, pH 6.5). The washed cells ($1\times10^7$ cells/ml) were then deposited onto 1x KK2/agar plates and allowed to develop over a 24-hour period, while images of the various multicellular structures were taken. Calcofluor staining (to stain cellulose and visualize fruiting body stalks and spores) was done as previously described[8], where individual fruiting bodies were picked using a forcep and stained with 0.01% Calcofluor white. Spores were also picked from sori for Calcofluor staining using a pipette tip. Images of Calcofluor stained structures were captured using a Zeiss LSM 710 confocal microscope.

*Spore viability*

To assess spore viability and germination, we followed a previously described protocol[9]. $2.5\times10^{6}$ cells were plated on 1 cm nitrocellulose filters on 1x KK2 agar to develop into fruiting bodies. These fruiting bodies were then collected and placed in 1x KK2 wash buffer and vortexed with 0.1 % Triton-X100. The total number of spores was also quantified at this time to determine the number of cells undergoing sporulation (measure of sporulation). The detergent resistant spores were clonally plated on lawns of bacteria, and then after 2-5 days, the resulting plaques for the survival of a single spore were counted (measure of germination).

*Cyclic di-GMP quantification*

Samples for cyclic di-GMP are derived from cells that were submerged in 1x KK2 buffer and starved for 12 hours until mound formation occurred. The supernatant was then collected and directly used for quantification of the secreted cyclic di-GMP fraction. Then, mounds were disassociated, and cells were lysed for 30 minutes in M-PER lysis buffer (ThermoFisher Scientific) on ice. The lysate was then used for the analysis of intracellular di-GMP. Cyclic di-GMP levels were then measured for samples using an ELISA kit (Cayman Chemical) according to the manufacturer’s instructions.

*RNA extraction and RT-qPCR*

For gene expression analyses on developing *Dictyostelium* cells, we collected multicellular structures at the indicated hours since the plating of cells (16, 18, 20, 22, 24 and 26) from slug formation (16h) to fruiting body formation (26h). These structures were then vortexed and dissociated before incubation in Trizol reagent (Thermofisher Scientific) according to the manufacturer's directions. For the ESCRT screen, we screened 30 characterized ESCRT genes from ESCRT-0, ESCRT-I, ESCRT-II, ESCRT-III, and ESCRT-IV[10]. shRNAs targeting PRP4K were induced using doxycycline in MCF7 cells and harvested after 72 hours for RNA. For human cells, the cells were lysed and homogenized with Trizol reagent before being used for RNA extractions. Samples were all processed while fresh for RNA extraction.

RNA was isolated using the Ambion PureLink RNA Mini Kit (Thermofisher Scientific) according to the manufacturer's protocol, with the addition of an on-column DNase I digestion. The quality of RNA was measured using a Nanodrop 2000 spectrophotometer (Thermofisher Scientific). Absorbance measurements A260/A280 and A260/A230 with ratios ~2.0 were accepted as pure for RNA. One microgram of RNA was reverse-transcribed to cDNA using the BioRad 5X iScript RT supermix kit (BioRad Laboratories Canada; Mississauga, ON, CA) for Reverse transcriptase quantitative PCR (qPCR), after which samples were diluted 1:1 with nuclease-free water. Reverse transcriptase lacking samples were included as a control to confirm no genomic DNA contamination.

RT-qPCR was performed on both *Dictyostelium* and human cDNA samples using the 2X SsoAdvanced Universal SYBR Green Supermix (BioRad). The reactions and all experiments were done in triplicate using a BioRad CFX Connect. NCBI Primer Blast (https://www.ncbi.nlm.nih.gov/tools/primer-blast/) was used to design all the primers (Supplementary Table 1). Gene expression data were normalized to at least two reference genes for *Dictyostelium* and human samples as indicated and analyzed using the BioRad CFX Maestro Software. Data were collected and analyzed as per the MIQE guidelines[11].

*Splicing assays*

Changes in splicing of transcripts for target gene *CHMP4B* was assessed using an RT-qPCR approach. Two sets of primers were generated: one set targeting the UTR and Exon 1 region of *CHMP4B* and the other set targeting exon junctions. The former will amplify all transcripts while the latter will only amplify transcripts that have undergone splicing. The ratio of these two sets of primers provides an index of relative splicing in the presence or absence of PRP4K. qPCR reactions were conducted as aforementioned using the two different sets of primers. A similar approach was taken to assess splicing of *vps32* in *Dictyostelium*. Primers are listed in Supplementary Table 1.

*siRNA and shRNA knockdown of PRP4K in human cells*

Human PRPF4B ON-TARGETplus siRNA oligos were purchased from Dharmacon (Dharmacon, Lafayette, CO, USA). Prior to transfection, MCF7 cells were seeded at 2.4 x $10^6$ cells/mL into 6-well or 10 cm plates overnight. Twenty-four hours after, MCF7 cells were transfected with 100 µM PRP4K siRNA or control siRNA. To achieve optimal transient knockdown, cells were incubated at 37°C with 5% $CO_2$ for 72 hours for protein analysis. To knockdown PRP4K in the HeLa cell lines using a dox-induction model, PRP4K-targeting TRIPZ Inducible Lentiviral shRNAs (shPRP4K-1 = clone: V2THS_47787, shPRP4K-2 = clone: V3THS_383962, shPRP4K-3 = clone: V3THS_383960) were purchased from Thermo Scientific. To create the MCF7 PRP4K knockdown cell lines, PRP4K-targeting TRIPZ Inducible Lentiviral shRNAs (Thermos Scientific) (shPRP4K-1 = clone: V2THS_47787, shPRP4K-2 = clone: V3THS_383960) were purchased. To select for viral integration, cells were cultured in 1 µg/mL puromycin for at least 48 h. To induce expression of the inducible TRIPZ PRP4K shRNA, 2 μg/mL doxycycline was added to culture media for 72 h with the drug being replaced every 24 h.

*Immunostaining and Immunofluorescence*

Cells were cultured and plated onto sterile coverslips (Fisher Scientific, catalog# 12-541-B) in a 6-well plate overnight. Cells were then washed with PBS and fixed in 4% paraformaldehyde for 30 minutes at room temperature. Coverslips were then rinsed in PBS and permeabilized using 0.5% Triton X-100 (Sigma-Aldrich, catalog# T8787) for 10 minutes at room temperature. In a humidity chamber, cells were blocked in PBS/4% bovine serum albumin (BSA) for 30 minutes and then immediately placed in primary antibody (1:200-1000) that has been diluted in PBS/4% BSA for one hour at room temperature. Primary antibodies include anti-p62 (Cell signalling technologies; Cat#7695), anti-p80 (Developmental Studies Hybridoma Bank; Cat#H161) and anti-FLAG (Sigma; Cat#F1804). Coverslips were then rinsed 3 times in PBS and incubated with an Alexa Fluor secondary antibody (Life Technologies) diluted in PBS/4% BSA for 45 minutes in the dark. Following the secondary antibody incubation, cells were washed 3 times (5 minutes per wash) with PBS and DAPI (Sigma-Aldrich, D9564) diluted at 1:1000 (1 $\mu$g/mL) was added to the second wash to visualize nuclei. Coverslips were mounted on sterile frosted glass microscope slides (Fisher Scientific, 12-550-15) using Dako fluorescent mounting medium (Dako, 106822-003). Immunofluorescence images were captured using a Zeiss LSM 710 confocal microscope under a 40X or 63X immersion oil objective lens. Images were processed using only linear adjustments (e.g., brightness/contrast) with Slidebook (Intelligent Imaging Innovations, Boulder, CO) and Adobe Photoshop 2020 (Version 22.0.1).

*mCherry-GFP-LC3 flux reporter assay*

The reporter cell line was generated using lentivirus encoding pBABE-puro-mCherry-GFP-LC3B (Addgene #22418)[12]. To produce lentivirus, HEK293T cells were plated on 10 cm tissue culture dishes and grown to 60-70% confluency. The cells were then co-transfected with 4.5 µg of pBABE-puro-mCherry-GFP-LC3B and 4 µg of pHIT and 0.5 µg of L-VSVG mixed with PEI, which were used as packaging vectors. After 48 hours, the HEK293T media was collected and filtered using a 0.45µ polystyrene filter and stored at 4°C. The virus was tittered by transducing HeLa and MCF7 cells using dilutions of the virus stock, which was followed by a media change after 24 hours and selected using 1 µg/ml puromycin.

*Visualization of endocytic trafficking*

For visualization of endocytic compartments, we incubated live HeLa cells with Vybrant™ DiO Cell-Labeling Solution (ThermoFisher Scientific) to visualize the endocytic pathway in cells. Cells were starved overnight in DMEM media (Sigma) containing 0.5% FBS (Wisent), and then incubated with fresh media containing 10% FBS and the DiO cell-labeling solution. The initial labelling of DiO was done on ice to prevent/slow internalization. Cells were also co-labeled with LysoTracker Red DND-99 (ThermoFisher Scientific) at 37 degrees for 1 hour.

*Transmission electron microscopy (TEM)*

Human cells were harvested and fixed for a minimum of 2 hours with 2.5% glutaraldehyde (pH 7.2) that was diluted with 0.1 M of sodium cacodylate buffer. After 2 hours, the fixed cells were rinsed 3 times (10 minutes per wash) with 0.1 M sodium cacodylate buffer.

The supernatant was removed by aspiration, and the cells were post-fixed for 2 hours in 1% osmium tetroxide. The osmium-fixed preparation was rinsed in distilled water and placed in 0.25% uranyl acetate at 4°C overnight. The following day, the fixed cells were dehydrated in a graded aqueous-acetone series, and infiltrated with, and embedded in, 100% epon-araldite resin in a 60°C oven for 48 hours to harden properly. Ultrathin sections were obtained using a Reichert Jung Ultracut E Ultramicrotome fitted with a diamond knife (approximately 100 nm thick). Sections were collected on 300 uncoated copper mesh grids, stained with 2% aqueous uranyl acetate for 10 minutes and then rinsed 2 times (5 minutes per wash) with distilled water. Sections were then stained with lead citrate for 4 minutes and rinsed 2 times (5 minutes per wash) with distilled water and observed using a JEOL JEM 1230 transmission electron microscope at 80 kV. Images were captured using a Hamamatsu ORCA-HR digital camera.

*Western blotting*

*Dictyostelium* cells were harvested during growth and washed in KK2 and lysed at 4°C in ice-cold lysis buffer (0.5% NP-40 in KK2 and 1X protease inhibitor [PMSF and P8340]) for 30 minutes. For human cell lines, the cells were harvested and washed with cold 10X PBS (pH. 7.4) and lysed at 4°C in ice-cold lysis buffer (0.5M Tris-HCl, pH. 7.4, 1.5M NaCl, 2.5% deoxycholic acid, 10% NP-40, 10mM EDTA, and 1X protease inhibitors [PMSF and P8340]) for 30 minutes on ice. Cell lysates were then pelleted by centrifugation at 16,000 g for 20 minutes in a 4°C precooled centrifuge. Lysates were then mixed 1:1 in 2X sample buffer (4% SDS, 20% glycerol, 10% 2-mercaptoethanol, 0.004% bromophenol blue, 0.125 M Tris HCl pH. 6.8) and boiled for 5 minutes prior to separation by SDS-PAGE.

Protein concentrations were determined using Bio-Rad Protein Reagent (Bio-Rad, 500-0006). Equal amounts of total protein were loaded and were run on precast gradient SDS-PAGE gels (Bio-Rad) at 50 V for 30 minutes and then 130 V for two hours. After electrophoresis, proteins were transferred to a 0.2 µm nitrocellulose membrane or PVDF membrane (activated in 100% methanol for 1 minute and then equilibrated in transfer buffer for 10 minutes) at 100 V for one hour. Membranes were then blocked in TBST/5% milk for one hour to minimize non-specific binding. Primary antibody staining (1:1000- 1:2500) occurred overnight at 4°C in TBST/2% BSA.

Primary antibodies used for western blotting include anti-PRPF4B (Novus Biologicals; Cat#NBP1-82999), Anti-CHMP4A antibody (Abcam; Cat#ab67058), anti-CHMP4B antibody (Abcam; Cat#ab135154), Anti-CHMP4C antibody (Abcam; Cat#ab168205), anti-LC3A/B (Abcam; Cat#ab58610), anti-P62 (Cell signalling technologies; Cat#8025), anti-FLAG (Sigma; Cat#F1804), and anti-GFP (Abcam; Cat#ab1218). The *Dictyostelium* loading controls, anti-β-actin (Cat#224−236−1), anti-α-tubulin (Cat#12G10), and anti-α-actinin (Cat#47−18−9), were purchased from the Developmental Studies Hybridoma Bank (University of Iowa, Iowa City, Iowa, USA). Rabbit polyclonal anti-Atg8b was provided as a gift by Dr. Ludwig Eichinger (University of Cologne). We also generated and used a polyclonal antibody against *Dictyostelium* Prp4k, which was raised in rabbits against a synthetic peptide corresponding to the epitope 12SVNDDKTNHGENLTC26 and then affinity purified (Genscript, Piscataway, New Jersey). The antibody was validated with our CRISPR/Cas9 KO *prp4k* cell lines (Figure S1). The following day, primary antibodies were detected using appropriate HRP-conjugated secondary antibody (1:5000) and visualized using ECL substrate (Bio-Rad, 1705061), which is detected by the ChemiDoc Imaging System (Bio-Rad) or film. Densitometric analyses of immunoblots were performed using ImageJ (NIH). All uncropped blots used in Figures are included in supplemental data (Supplementary Figure 9 and Supplementary Figure 10).

*Plasmid construction*

Plasmids for the expression of *Dictyostelium* proteins Prp4k and Vps32 were generated using the total RNA collected from 2 x $10^7$ cells starved on petri dishes while submerged in KK2 for 4 hours (where expression of *prp4k* and *vps32* is high). These samples were then processed for total RNA as described previously for RT-qPCR using Trizol reagent (Thermofisher scientific) and the Ambion PureLink RNA Mini Kit (Thermofisher Scientific). The cDNAs for *prp4k* and *vps32* were amplified using the SuperScript™ III One-Step RT-PCR System with Platinum™ *Taq* DNA Polymerase (Thermofisher Scientific). The cDNAs were then cloned by restriction digest cloning (BamHI and SalI) into the actin 15 (*act15*) promoter with N-terminally tagged FLAG expression vector, pTX-FLAG (digested with BamHI and XhoI). cDNA for human CHMP4B was obtained from ThermoFisher and used for cloning into the 3xFLAG-J1 vector by restriction digest cloning (BamHI and SalI).

*Statistics*

The graphs in the study were generated using GraphPad Prism 9. For statistical analyses between groups of 3 or more, significance was determined using a One-way ANOVA, with Tukey's post-hoc analysis used for comparison. The exception being with the data from RT-qPCR analyses on *Dictyostelium development* at multiple timepoints, where a Two-way ANOVA, with Dunnett's post-hoc analysis used for comparison. For statistical analyses between groups of 2 (siPRP4K experiments), a two-tailed Student's t-test was used for comparisons. All statistical analyses were done using GraphPad Prism 9.
